# Combinatorial Wnt signaling landscape during brachiopod anteroposterior patterning

**DOI:** 10.1101/2023.09.03.556047

**Authors:** Bruno C. Vellutini, José M. Martín-Durán, Aina Børve, Andreas Hejnol

## Abstract

**Background:** Wnt signaling pathways play crucial roles in animal development. They establish embryonic axes, specify cell fates, and regulate tissue morphogenesis from the early embryo to organogenesis. It is becoming increasingly recognized that these distinct developmental outcomes depend upon dynamic interactions between multiple ligands, receptors, antagonists, and other pathway modulators, consolidating the view that a combinatorial “code” controls the output of Wnt signaling. However, due to the lack of comprehensive analyses of Wnt components in several animal groups, it remains unclear if specific combinations always give rise to specific outcomes, and if these combinatorial patterns are conserved throughout evolution.

**Results:** In this work, we investigate the combinatorial expression of Wnt signaling components during the axial patterning of the brachiopod *Terebratalia transversa*. We find that *T. transversa* has a conserved repertoire of ligands, receptors, and antagonists. These genes are expressed throughout embryogenesis but undergo significant upregulation during axial elongation. At this stage, Frizzled domains occupy broad regions across the body while Wnt domains are narrower and distributed in partially overlapping patches; antagonists are mostly restricted to the anterior end. Based on their combinatorial expression, we identify a series of unique transcriptional subregions along the anteroposterior axis that coincide with the different morphological subdivisions of the brachiopod larval body. When comparing these data across the animal phylogeny, we find that the expression of Frizzled genes is relatively conserved, whereas the expression of Wnt genes is more variable.

**Conclusions:** Our results suggest that the differential activation of Wnt signaling pathways may play a role in regionalizing the anteroposterior axis of brachiopod larvae. More generally, our analyses suggest that changes in the receptor context of Wnt ligands may act as a mechanism for the evolution and diversification of the metazoan body axis.

## Background

Wnt genes play multiple roles during embryogenesis [1, 2]. They encode secreted glycoproteins with a conserved series of cysteine residues that often act as a symmetry-breaking signal [3]. Wnt activity can establish embryonic axes [4, 5], mediate cell fate decisions in early embryos [6–8], and specify endomesodermal tissues during gastrulation [9–13]. Moreover, they can also control morphogenetic processes, such as apical constriction, convergent extension, and cell migration [14–20]. This multitude of roles is consistent with the sheer complexity of Wnt signaling pathways, which involves 13 subfamilies of Wnt ligands, five subfamilies of Frizzled receptors, additional co-receptors, and different agonists, antagonists, downstream players, and effector molecules. In general terms, there are at least three interconnected Wnt pathways. The Wnt/beta-catenin (canonical) pathway regulates cell fate specification through the activity of beta-catenin, the Wnt/PCP (planar cell polarity) pathway controls cell polarity during tissue morphogenesis, and the Wnt/calcium pathway regulates intracellular calcium levels for convergent extension movements—although these functions are not exclusive, as the Wnt/PCP and Wnt/calcium pathways can also control cell fate specification through the activity of other transcriptional effectors like ATF2 and NFAT, respectively [21, 22]. Understanding how these intricate signaling networks regulate embryogenesis and influences developmental evolution remains a significant challenge.

The discovery of staggered Wnt expression domains in the tail bud of amphioxus embryos [23] and along the body axis of sea anemone embryos [24] raised the hypothesis that different combinations of Wnt genes can pattern different body regions. This idea, commonly referred to as the *Wnt code* or *landscape* [25–27], is an analogy to the *Hox code*, where the combinatorial expression of nested Hox domains determines the positional identities of tissues along the body axis [28]. Over the years, however, it has been increasingly recognized that the output of Wnt signaling does not depend solely on the expression of Wnt ligands. In fact, Wnt pathways operate via an intricate network of dynamic protein interactions where the downstream response depends on the local availability of receptors, the presence of different antagonists, and the activity of pathway modulators [29, 30]. That means that, depending on the receptors present in the tissue, one Wnt ligand can activate or inhibit a different Wnt pathway and thus determine processes as diverse as fate specification, cellular organization, and tissue morphogenesis [31–33]. This complexity makes the Wnt code particularly challenging to elucidate.

A necessary step to untangle this combinatorial code is to extend the analyses of ligand–receptor contexts of Wnt genes to other animal groups using a comparative approach [34]. The comparison between flies and other animals was crucial to reveal the broader importance of the Hox genes as a high-level axial patterning system and not merely an arthropod-specific feature linked to body segmentation [35–37]. But while the expression of some Wnt genes is conserved in different animals (e.g., *wnt8* is expressed in the neuroectoderm of vertebrates [38, 39] and in spiders [27], annelids [40], and hemichordates [41]), it remains unclear if the combinatorial expression of Wnt signaling components along the embryonic body axis is conserved across the animal phylogeny.

The developmental Wnt landscape has been widely studied in several animal groups, but fewer works have analyzed the receptor context information in the Spiralia, a major branch of bilaterian animals with diverse body patterns [42, 43]. While ecdysozoans have lost several Wnt genes [44–46], spiralians have retained the ancestral Wnt complement [27, 47, 48], indicating that they can be an informative group to understand the role of Wnt signaling in axial patterning evolution. However, most analyses about Wnt genes in Spiralia were performed in annelids and mollusks, and the expression data in other spiralian lineages is still lacking.

Brachiopods are sessile spiralians with bivalve shells [49]. They have a reduced adult morphology, but complex embryogenesis where a radially symmetric gastrula undergoes a series of morphogenetic changes to form a larval body subdivided into a series of distinct lobes along the anteroposterior axis [50, 51]. In previous studies using the rhynchonelliform brachiopod *Terebratalia transversa* (Sowerby, 1846)—a species with a well-characterized embryonic development [52–62]—we found that Wnt signaling plays a role in the specification of the endomesoderm and posterior fates [63], and in the patterning of the head–trunk boundary [64]. Moreover, we found that over-activation of the Wnt/beta-catenin pathway disrupts the molecular and morphological organization of the larval subdivisions [63, 64], suggesting that Wnt activity contributes to the axial patterning of the larva. However, a full characterization of the Wnt signaling components and their developmental expression, including the ligand–receptor contexts is lacking for *T. transversa*, and for brachiopods in general.

In this study, we characterize the Wnt complement of the brachiopod *T. transversa* and investigate the spatiotemporal expression of Wnt signaling components throughout embryogenesis. We find that during axial elongation, the expression of ligands, receptors, and antagonists show an anteroposterior organization forming regionalized transcriptional territories, each expressing a unique combination of transcripts. These territories precede and coincide with the morphological subdivisions of the larval body, indicating that the differential activation of Wnt signaling may contribute to pattern the brachiopod larval body. We identified differences in receptor-context that may be involved in patterning an evolutionary novelty of lecithotrophic brachiopod larvae, the reversible mantle lobe. A comparative analysis reveals that while the expression of Frizzled receptors is evolutionarily conserved, the expression of Wnt ligands is more variable. This suggests that the evolutionary shuffling in the expression of Wnt ligands may be a mechanism underlying the evolution of anteroposterior diversification in bilaterians.

## Results

### *Terebratalia transversa* has a conserved repertoire of Wnt genes

Metazoans have a large Wnt repertoire with 13 subfamilies [24, 48, 65]. To characterize the Wnt complement of the brachiopod *Terebratalia transversa*, we surveyed a reference transcriptome of the species for Wnt genes using similarity searches with known Wnt genes from other animals. We identified 13 Wnt genes with representatives of 12 of the 13 Wnt subfamilies (Fig. 1). *T. transversa* is missing *wnt3*, a gene known to have been lost in Protostomia [27, 48], and has two copies of *wnt1*. One of the *wnt1* paralogs—named hereafter *wnt1t*—has a conserved Wnt domain, but is highly divergent at the sequence level compared to other *wnt1* orthologs across bilaterians (<u>Additional file 1: Fig. S1</u>). Our phylogenetic analysis suggests that this paralog originated via a lineage-specific duplication within *T. transversa* or rhynchonelliform brachiopods (<u>Additional file 2: Fig. S2</u>). Besides the loss of *wnt3* and duplication of *wnt1*, *T. transversa* shows a single representative ortholog for the remaining subfamilies, suggesting that the ancestral repertoire of metazoan Wnt genes remained largely conserved.

**Figure 1:**
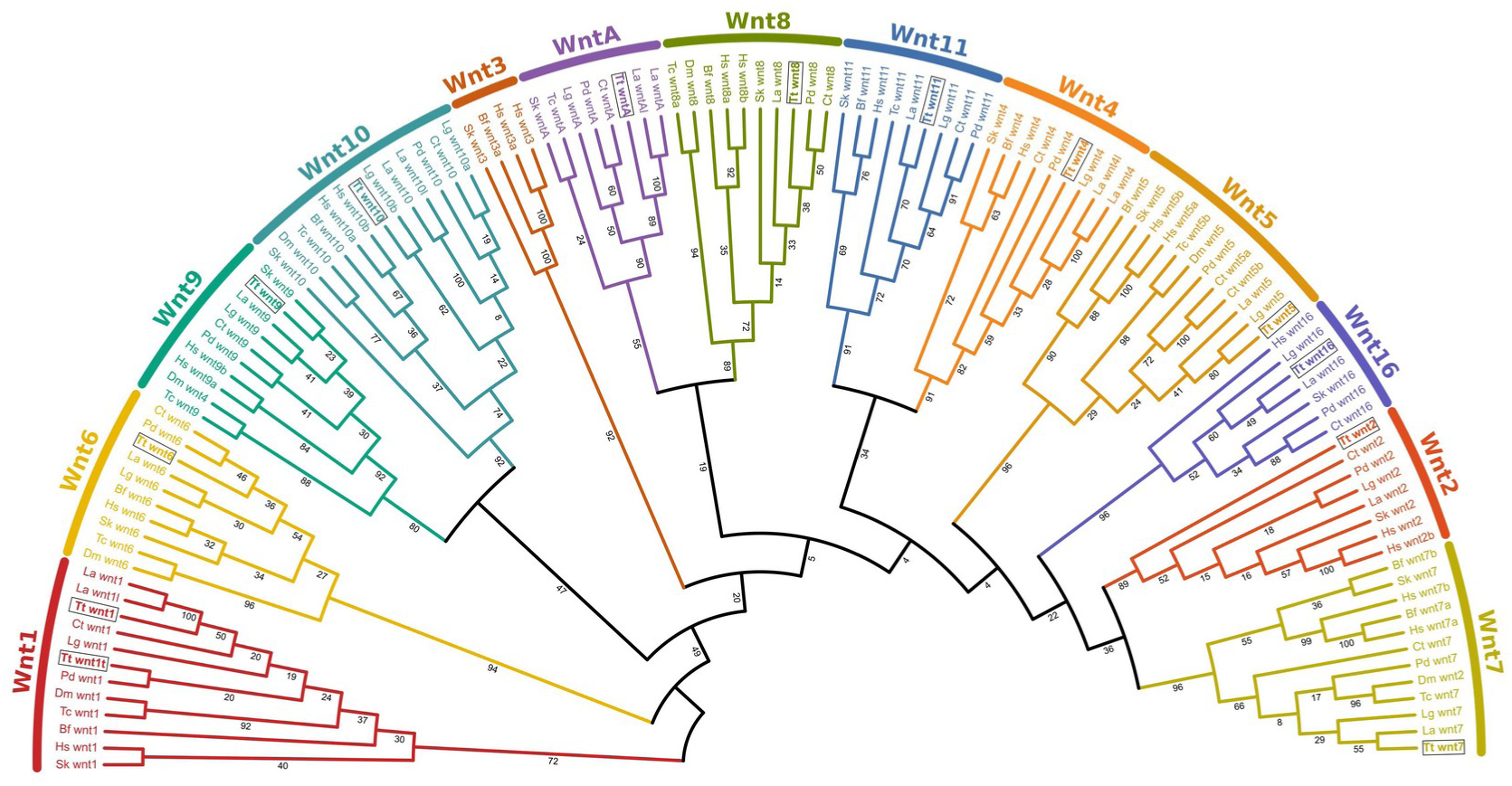
Orthology assignment of *Terebratalia transversa* Wnt genes. Best-scoring tree of a maximum likelihood phylogenetic analysis using the amino acid sequences of known metazoan Wnt genes. The color-coding represents different Wnt subfamilies and the numbers show the support values of individual branches. *Terebratalia transversa* (Tt) orthologs are outlined by a box. The other species are *Branchiostoma floridae* (Bf), *Capitella teleta* (Ct), *Drosophila melanogaster* (Dm), *Homo sapiens* (Hs), *Lingula anatina* (La), *Lottia gigantea* (Lg), *Platynereis dumerilii* (Pd), *Saccoglossus kowalevskii* (Sk), and *Tribolium castaneum* (Tc).

### Wnt genes are upregulated in concert during axial elongation

To uncover the developmental dynamics of Wnt expression in *T. transversa*, we analyzed stage-specific RNA-Seq data from the unfertilized egg to the post-metamorphic juveniles (Fig. 2 and Table 1). We detect no Wnt transcripts expressed in the oocyte or mid blastula stages (the high levels of *wnt4* and *wntA* in early stages is due to a bias in the expression quantification, see Methods for a detailed explanation). Wnt expression shifts significantly at the late blastula stage (19h), when a concerted upregulation of *wnt1*, *wnt1t*, *wnt8*, *wnt10*, and *wnt16* occurs (Fig. 2). Throughout gastrulation, Wnt genes continue to be upregulated with *wnt1* and *wnt5* in the early gastrula (26h); *wnt6*, *wnt7*, and *wnt11* in the mid gastrula (37h); and *wnt2*, *wnt9*, and *wnt10* in the late gastrula (51h). Between the late gastrula and early larva, all Wnt genes are expressed, but some are downregulated after gastrulation (*wnt6* and *wnt10*) and after metamorphosis (*wnt7* and *wnt16*) (Fig. 2). Therefore, Wnt expression is dynamic throughout development but peaks late in gastrulation, when the body elongates along the anteroposterior axis, and at the onset of the morphological differentiation of the larval lobes in *T. transversa*.

**Figure 2:**
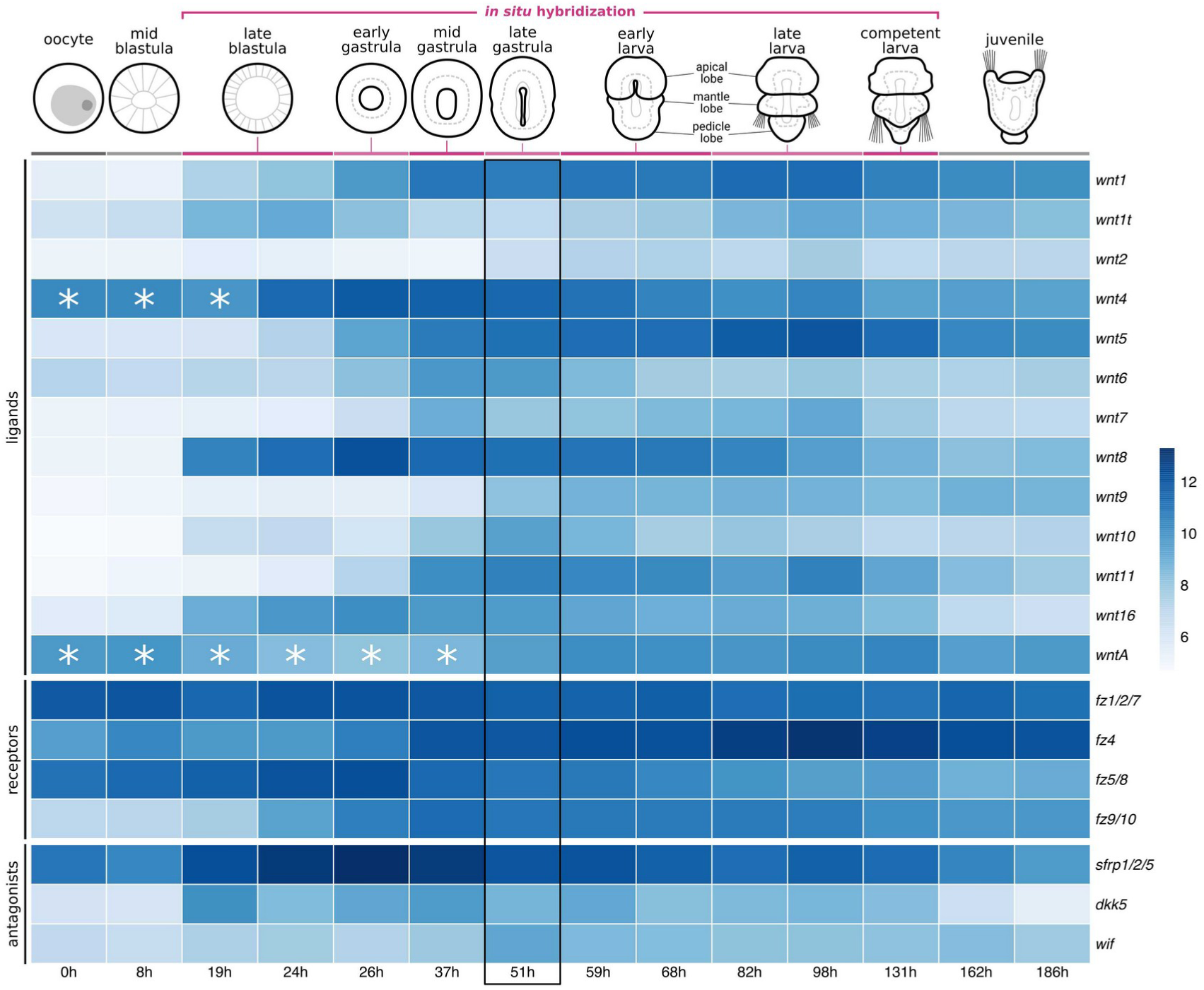
Expression of Wnt signaling components during *Terebratalia transversa* development. The heatmap represents the log-normalized transcript counts for ligands, receptors, and antagonists calculated from stage-specific RNA-Seq data. Each cell shows the average value between two replicates. Asterisks in *wnt4* and *wntA* denote samples where the expression levels were overestimated due to the expression of an antisense gene present in the same transcript (see Methods for details). The black outline marks the late gastrula stage (51h), when all Wnt genes are expressed. The illustrations depict *T. transversa* developmental stages from the oocyte until the postmetamorphic juvenile. The stages we analyzed using *in situ* hybridization (early gastrula to late larva) are highlighted in magenta.

**Table 1:**
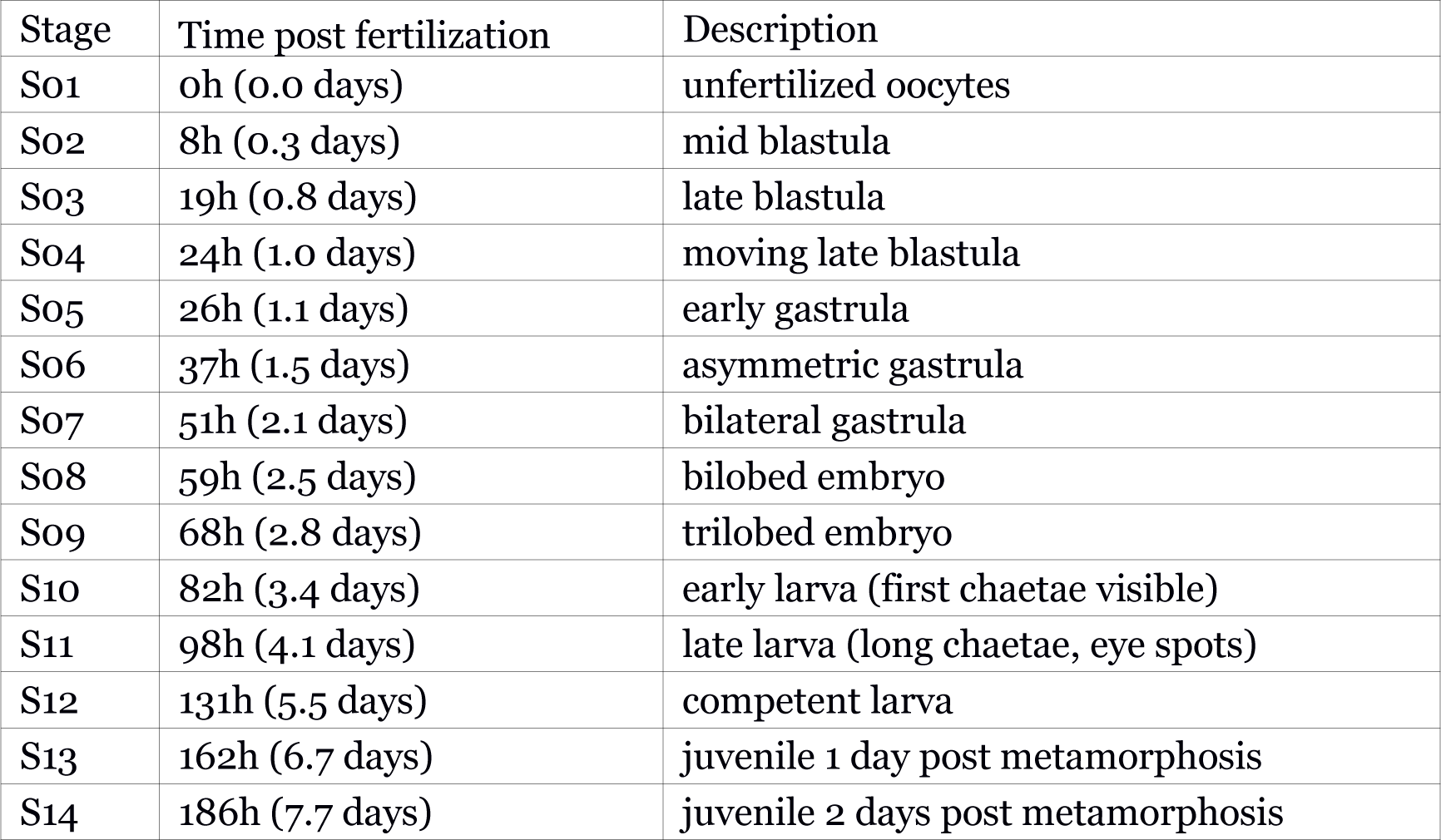
*Developmental stages sampled for the stage-specific transcriptome of Terebratalia transversa*.

### Wnt expression domains partially overlap along the anteroposterior axis

To uncover the spatial localization of Wnt ligands during brachiopod development, we performed *in situ* hybridization in *T. transversa* embryos from late blastula to competent larva (Fig. 3, 4, 5, <u>Additional file 3: Fig. S3</u>, <u>Additional file 4: Fig. S4</u>, and <u>Additional file 5: Fig. S5</u>).

**Figure 3:**
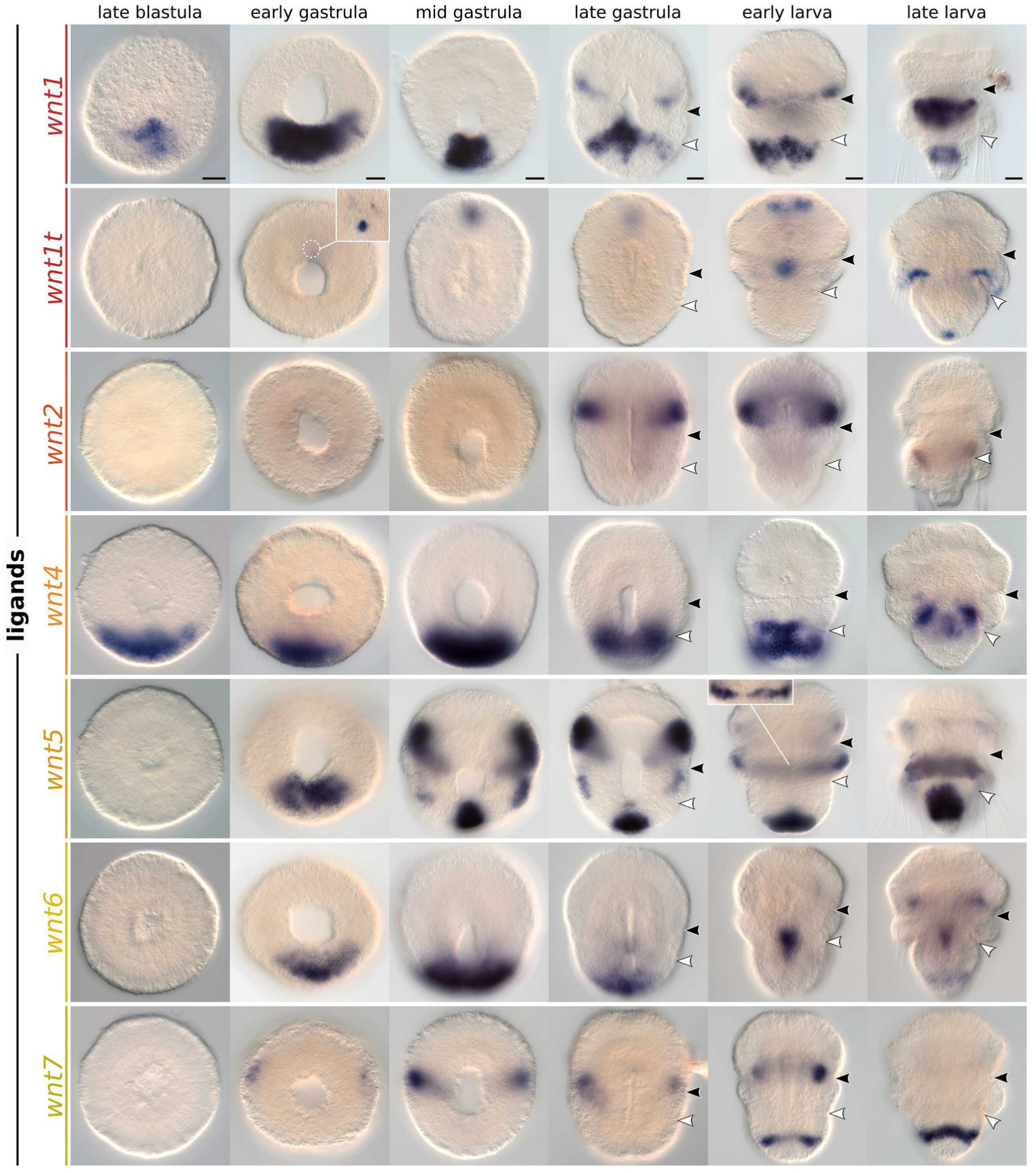
Whole-mount colorimetric *in situ* hybridization of *Terebratalia transversa wnt1*, *wnt1t*, *wnt2*, *wnt4*, *wnt5*, *wnt6*, and *wnt7*. The panels show representative expression patterns for the developmental stages between late blastula and late larva. The samples are oriented with a blastoporal/ventral view and anterior end to the top. Black arrowheads indicate the apical–mantle boundary. White arrowheads indicate the mantle–pedicle boundary. Panels for *wnt1* originally published under a Creative Commons Attribution License in [64] and reprinted here for completion. Scale bars = 20µm.

**Figure 4:**
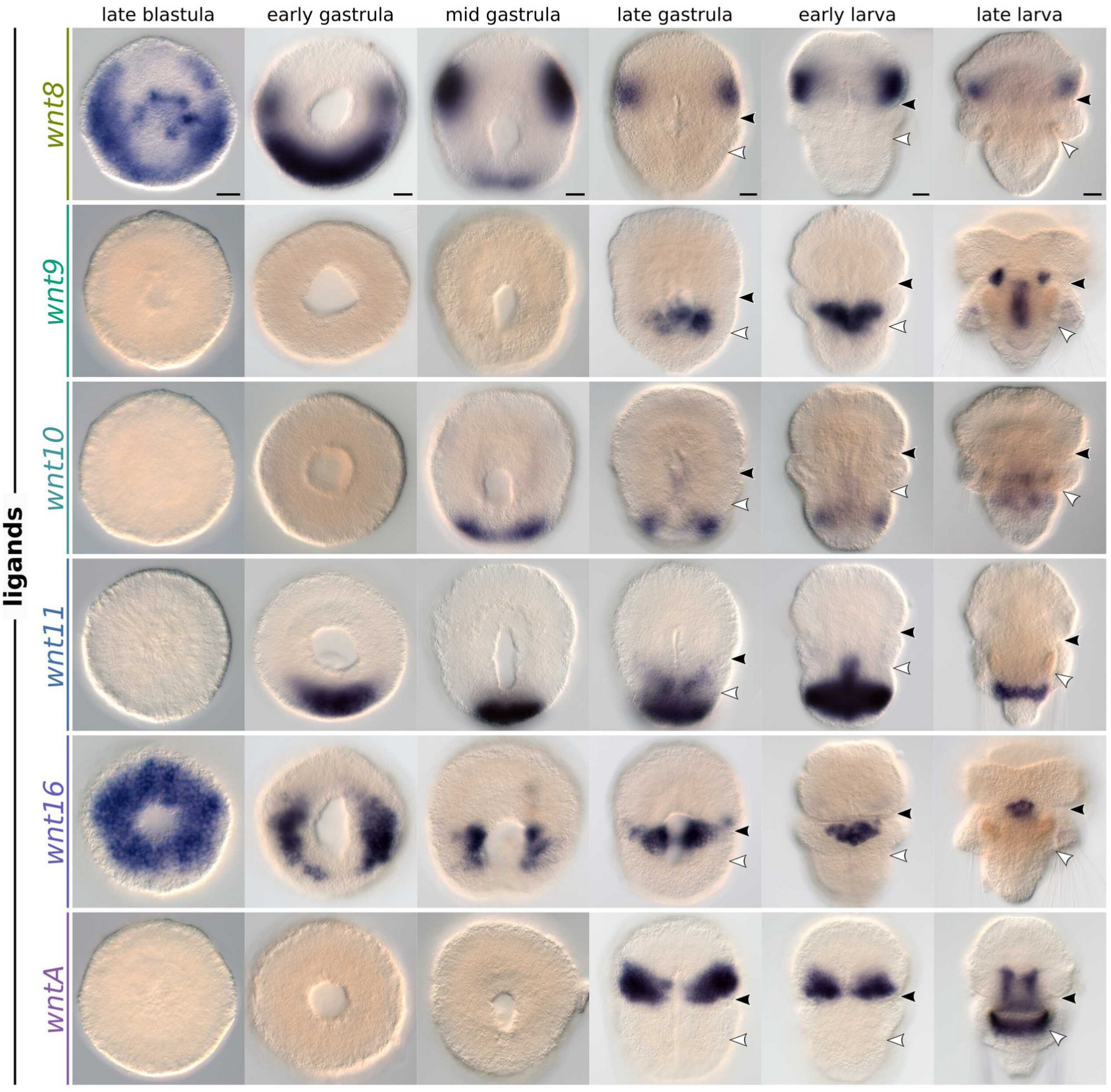
Whole-mount colorimetric *in situ* hybridization of *Terebratalia transversa wnt8*, *wnt9*, *wnt10*, *wnt11*, *wnt16*, and *wntA*. The panels show representative expression patterns for the developmental stages between late blastula and late larva. The samples are oriented with a blastoporal/ventral view and anterior end to the top. Black arrowheads indicate the apical–mantle boundary. White arrowheads demarcate the mantle–pedicle boundary. Scale bars = 20µm.

**Figure 5:**
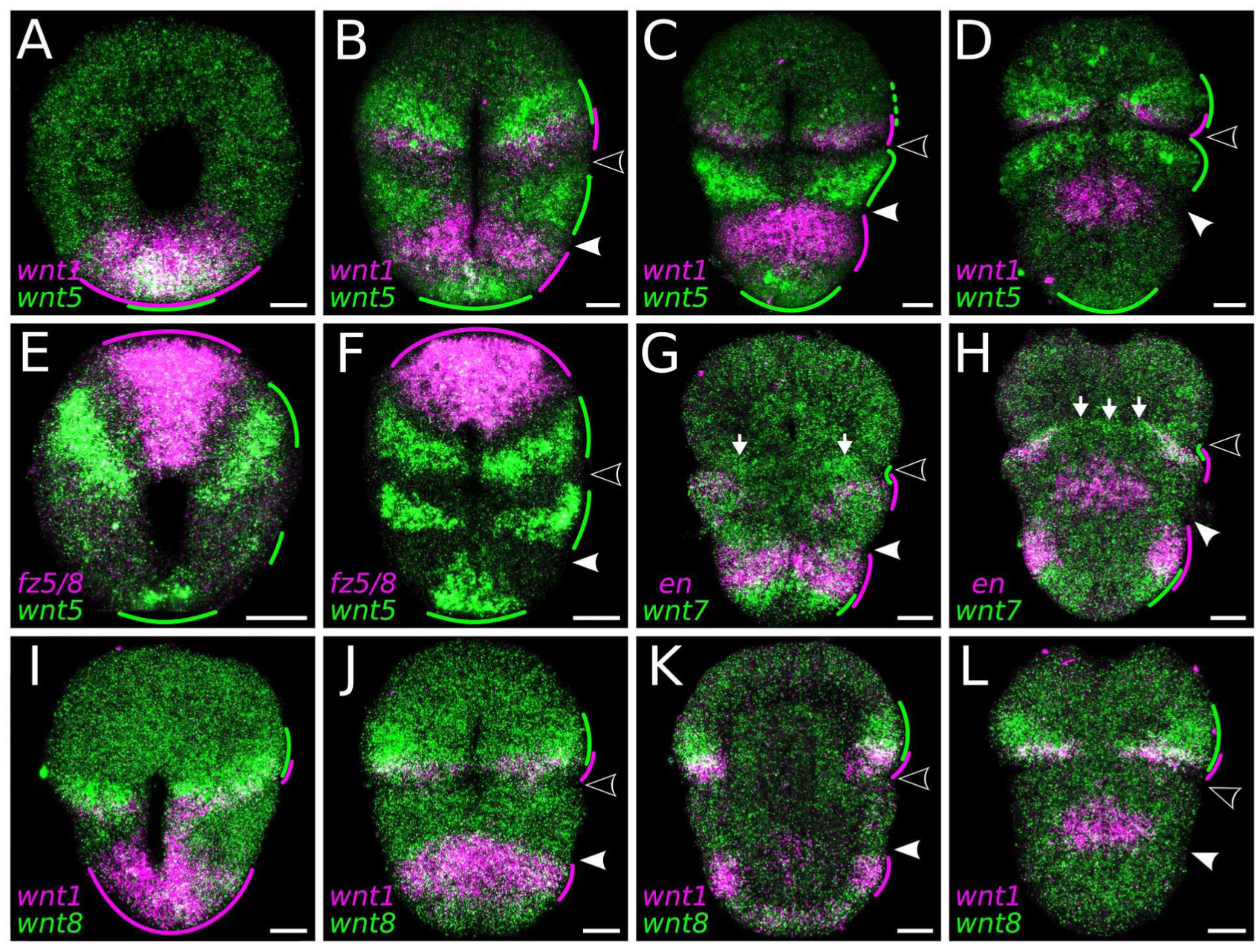
Whole-mount double-fluorescent *in situ* hybridization of *Terebratalia transversa* Wnt genes. (A–D) Expression of *wnt1* (magenta) and *wnt5* (green) in the early gastrula (A), late gastrula (B) and early larva in ventral (C) and dorsal (D) views. (E–F) Expression of *wnt5* (green) and *fz5/8* (magenta) in the mid gastrula (E) and late gastrula (F) in ventral views. (G–H) Expression of *wnt7* (green) *engrailed* (*en*) (magenta) in the early larva in ventral (G) and dorsal (H) views. (I–L) Expression of *wnt1* (magenta) and *wnt8* (green) in the mid gastrula (I) and late gastrula (J) in ventral views, late gastrula in an midbody optical section (K), and early larva in dorsal view (L). Green and magenta lines highlight the extension and overlap between domains. Areas in the tissue where the expression overlaps appear in white. Samples oriented with anterior end to the top. Black arrowheads indicate the apical–mantle boundary. White arrowheads demarcate the mantle–pedicle boundary. The panels show representative expression patterns for each sample. Scale bars = 20µm.

*wnt1* is expressed in the ectoderm and invaginating endomesoderm of the posterior blastopore lip in the late blastula (Fig. 3, <u>Additional file 3: Fig. S3</u>, and [64]). This domain expands laterally, forming a ventral ectodermal band at the anterior most portion of the pedicle lobe in the early larva, a region that gives rise to the ventral shell primordium in the late larva. From the late gastrula stage, *wnt1* is also expressed in a narrow ectodermal stripe around the posterior region of the apical lobe and in the dorsal shell primordium. The apical lobe expression fades, and a new *wnt1* domain appears encircling the posterior subdivision of the pedicle lobe in the late larva.

*wnt1t* expression domains differ significantly from its paralog *wnt1*. We first detect *wnt1t* transcripts in a single ectodermal spot at the animal pole of the early gastrula (Fig. 3 and <u>Additional file 3: Fig. S3</u>). This domain localizes to the anterior end of the embryo and is expressed until the early larva, when only subsets of cells continue expressing *wnt1t*. At this stage, a central patch of ventral ectoderm posterior to the mouth also begins expressing *wnt1t*. Finally, in the late larva, *wnt1t* is upregulated in an ectodermal ring beneath the mantle lobe, and at the terminal tip of the pedicle lobe ectoderm.

*wnt2* is only expressed in the late gastrula and early larva stages in bilateral ectodermal bands that encircle the posterior portion of the apical lobe almost entirely, except for the ventral and dorsal midlines (Fig. 3 and <u>Additional file 3: Fig. S3</u>).

*wnt4* is expressed at the posterodorsal ectoderm from late blastula to late gastrula (Fig. 3 and <u>Additional file 3: Fig. S3</u>). The pattern is similar to *wnt1*, but *wnt4* transcripts are localized more dorsally (<u>Additional file 5: Fig. S5A–C</u>). In the early larva, the expression at the posterior end and dorsal portion fades, the domain becomes narrower, and acquires a subterminal position within the ventral ectoderm of the pedicle lobe. This domain is still present in the late larva, when *wnt4* begins to be expressed in the posterior endoderm.

*wnt5* is expressed in three distinct ectodermal domains—in the apical, mantle, and pedicle lobes, respectively. We first detect expression in the early gastrula with transcripts at the posterior blastopore lip and anterolateral ectoderm (Fig. 3 and <u>Additional file 3: Fig. S3</u>). The posterior ectodermal domain is narrower than the *wnt1* domain (Fig. 3 and 5A) and maintains a terminal position until the late larva stage, when the tip of the pedicle lobe is cleared from expression (Fig. 3 and <u>Additional file 3: Fig. S3</u>). The anterolateral domains expand in the mid gastrula to encircle the posterior portion of the apical lobe ectoderm, and fade in the late larva. *wnt5* is also expressed in the leading edge of the growing mantle lobe ectoderm from mid gastrula to late larva. The ectodermal expression domains of *wnt5* and *wnt1* occupy distinct regions along the anteroposterior axis that coincide with the subdivisions of the larval lobes (Fig. 5B,C and <u>Additional file 5: Fig. S5G</u>).

*wnt6* transcripts localize to the posterior blastopore lip, similarly to *wnt1* and *wnt4*, from the early to the late gastrula (Fig. 3 and <u>Additional file 3: Fig. S3</u>). This ectodermal domain is cleared in the early larva, when *wnt6* is activated in a midbody section of the endoderm. In the late larva, we also detect *wnt6* domains in the ectoderm of the apical and pedicle lobes.

*wnt7* initiates as a lateral pair of anterior ectodermal stripes that progressively extend around the entire embryo circumference until the early larva (Fig. 3 and <u>Additional</u> <u>file 3: Fig. S3</u>). This *wnt7* stripe demarcates the apical–mantle boundary, partially overlapping with *wnt1*- and *engrailed*-expressing cells at the anteriormost region of the mantle lobe (Fig. 5G,H and <u>Additional file 5: Fig. S5H</u>; see also [64]). In the early larva, the anterior *wnt7* stripe disappears, and a posterior ectodermal stripe appears demarcating the boundary between the subterminal and terminal regions of the pedicle lobe, between the posterior territories of *wnt1* and *wnt5*.

*wnt8* is expressed in a ring of cells in the invaginating endomesoderm and in a broad ectodermal band encircling the late blastula (Fig. 4 and <u>Additional file 4: Fig. S4</u>). In the early and mid gastrula, *wnt8* transcripts are cleared from the endomesoderm and from the anterior ectoderm, and two distinct ectodermal domains remain: a pair of broad lateral territories in the apical lobe, and a wide posterodorsal domain in the pedicle lobe. The lateral territories expand ventrally and dorsally, encircling the apical lobe ectoderm, while the posterior ectodermal domain fades in the late gastrula. The anterior *wnt8* territories partially overlap with the anterior expression of *wnt1* in the apical lobe ectoderm (Fig. 5I–L and <u>Additional file 5: Fig. S5G</u>).

*wnt9* transcripts are first expressed in the invaginated endomesoderm of late gastrula embryos at a subterminal position (Fig. 4 and <u>Additional file 4: Fig. S4</u>). The domain forms a contiguous patch of mesodermal and endodermal cells expressing *wnt9* in the early larva, which differentiates into two distinct territories; one endodermal in the central portion of the gut and another mesodermal in a bilateral pair of anterior domains near the apical–mantle boundary.

*wnt10* is expressed from the mid gastrula stage in a posterior ectodermal domain, which acquires a subterminal position within the pedicle lobe in the early larva (Fig. 4 and <u>Additional file 4: Fig. S4</u>). Additionally, we detect *wnt10* transcripts in the late gastrula at a dorsal ectodermal patch of the apical lobe, similar to the dorsal domain of *wnt1t*, and in the late larva at the posterior mesoderm.

*wnt11* is expressed in a posterodorsal ectodermal domain in the early gastrula, similar to *wnt4* (Fig. 4 and <u>Additional file 4: Fig. S4</u>). The domain encircles the pedicle lobe ectoderm in the early larva, and becomes reduced to a narrow ectodermal stripe on the ventral portion of the pedicle lobe in the late larva. In the early larva, *wnt11* is also expressed in the ventral ectoderm at the mantle–pedicle boundary, and in the posterior endoderm of the larval gut (Fig. 4 and <u>Additional file 4: Fig. S4</u>).

*wnt16* is expressed in the invaginating endomesoderm and vegetal ectoderm around the blastopore in the late blastula (Fig. 4 and <u>Additional file 4: Fig. S4</u>). During gastrulation, the endomesodermal expression clears, and only the ectodermal domain remains as lateral patches near the blastopore lip. With the blastopore closure, *wnt16* forms a heart-shaped domain in the ectoderm and presumably mesoderm at the ventral midline of the mantle lobe in the early larva.

*wntA* appears in the mid gastrula as paired, ventral ectodermal domains located at the posterior portion of the apical lobe (Fig. 4 and <u>Additional file 4: Fig. S4</u>). In the late larva, these anterior ectodermal domains are cleared, and *wntA* expression is activated in paired, ventral mesodermal bands adjacent to the mouth.

Overall, Wnt genes are primarily expressed in the ectoderm, in diverse partially overlapping domains along the anteroposterior axis (<u>Additional file 6: Fig. S6</u> and <u>Additional file 7: Fig. S7</u>).

### Frizzled genes exhibit gradual expression changes throughout embryogenesis

Frizzled genes encode seven-pass transmembrane proteins with an extracellular cystein-rich domain and act as receptors in Wnt signaling pathways [66]. There are five Frizzled subfamilies in metazoans [67], but the subfamily *fz3/6* is only found in tunicates and vertebrates [68]. In the brachiopod *T. transversa*, we identified a total of four Frizzled genes with a single ortholog for the *fz1/2/7*, *fz5/8*, *fz9/10*, and *fz4* sub-families, respectively (<u>Additional file 8: Fig. S8</u>).

Frizzled receptors are expressed throughout *T. transversa* development. In the unfertilized oocyte, *fz1/2/7* and *fz5/8* are highly expressed (Fig. 2). The expression of *fz1/2/7* remains high from the oocyte to juvenile stages, while the expression of *fz5/8* peaks before gastrulation and decays over time. *fz4*, which is initially expressed at lower levels, peaks late in development, at the larval stages, an expression profile complementary to the one of *fz5/8* (Fig. 2). In contrast, *fz9/10* expression increases during gastrulation and remains relatively constant in subsequent stages.

Overall, each Frizzled shows a unique expression profile but in contrast to Wnt dynamic changes, the levels of Frizzled transcripts change more gradually during development.

### Frizzled expression domains occupy broad but distinct body regions

We carried out *in situ* hybridization for all Frizzled genes of *T. transversa* to reveal their domains of expression during axial elongation (Fig. 6).

**Figure 6:**
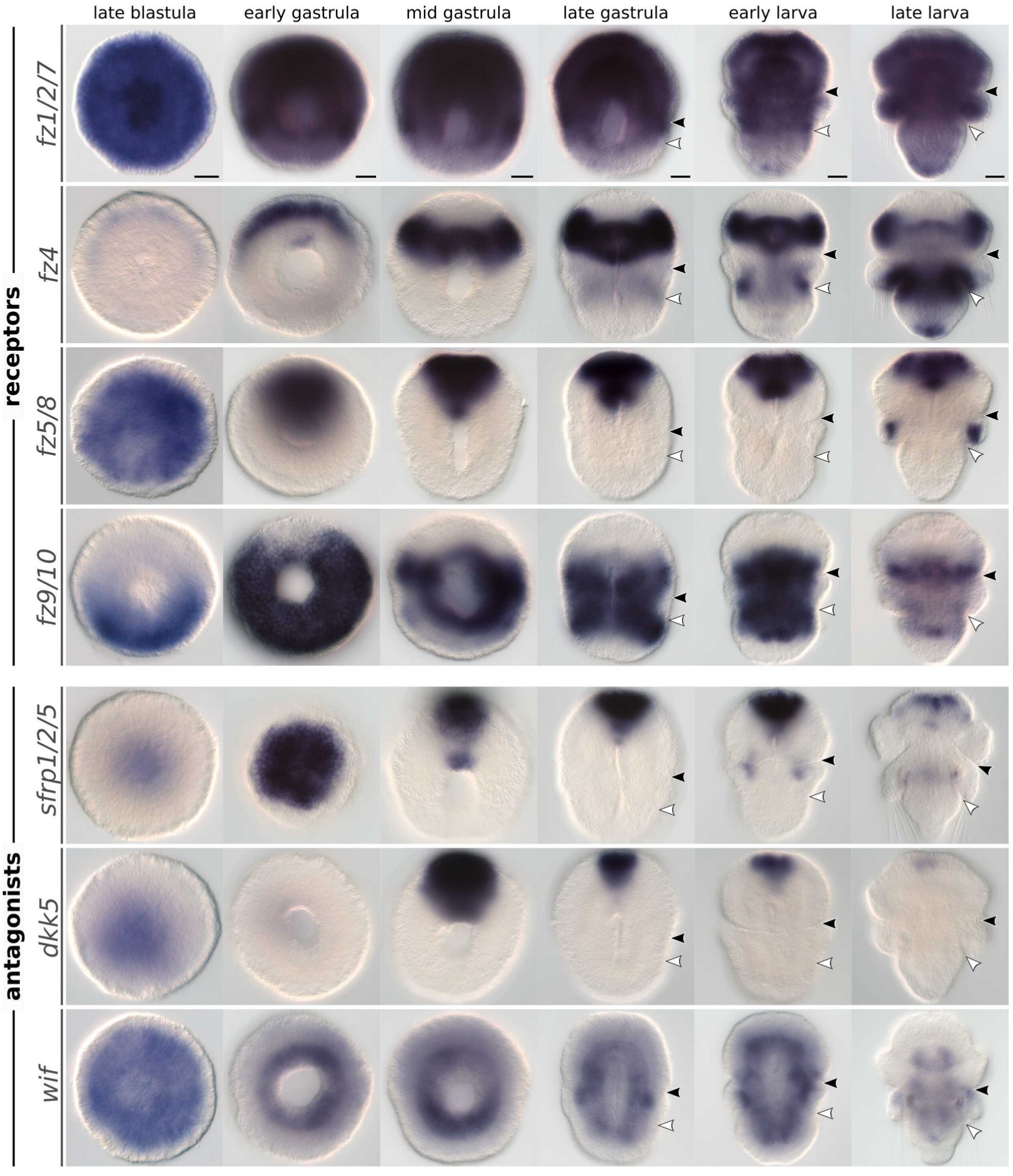
Whole-mount colorimetric *in situ* hybridization of *Terebratalia transversa* Frizzled and Wnt antagonist genes. Developmental stages between late blastula and late larva for *fz1/2/7*, *fz4*, *fz5/8*, *fz9/10*, *sfrp1/2/5*, *dkk5*, and *wif*. The panels show representative expression patterns for each sample. The samples are oriented with a blastoporal/ventral view and anterior end to the top, except for *sfrp1/2/5* early gastrula showing the animal pole. Black arrowheads indicate the apical–mantle boundary. White arrowheads demarcate the mantle–pedicle boundary. Scale bars = 20µm.

*fz1/2/7* expression is mostly ubiquitous (Fig. 6 and <u>Additional file 9: Fig. S9</u>). It is expressed in all tissues of the late blastula, with strong signal in the animal pole ectoderm and invaginating endomesoderm. During gastrulation, the anterior and middle portions of the body continue to express *fz1/2/7* across all germ layers, but the posterior transcripts get progressively cleared from the pedicle lobe tissues. In larval stages, *fz1/2/7* is up-regulated in the terminal portion of the pedicle lobe ectoderm, and becomes nearly ubiquitous again in the late larva.

*fz4* is first expressed in the animal pole ectoderm of the late blastula (Fig. 6 and <u>Additional file 9: Fig. S9</u>). These anterior transcripts form a subapical ectodermal ring encircling the animal pole of the early gastrula that localizes to the posterior portion of the apical lobe in subsequent stages. *fz4* is also expressed in the anterior mesoderm from the early gastrula. In the late gastrula, we detect *fz4* transcripts in the dorsal ectoderm between the mantle and pedicle lobes, a domain that becomes stronger in the late larva as it expands around the dorsal ectoderm of the pedicle lobe as well as in the infolded ectodermal and mesodermal tissues of the growing mantle lobe. Additionally, a *fz4* domain appears at the posterior tip of the pedicle lobe ectoderm in the late larva.

*fz5/8* is mainly expressed at the anteriormost region of the embryo’s ectoderm and mesoderm (Fig. 6 and <u>Additional file 9: Fig. S9</u>). We first detect transcripts in the animal pole ectoderm of the late blastula, these transcripts become restricted to the anterodorsal portion of the ectoderm in the early gastrula, and finally expand to the anteroventral ectoderm from mid gastrula to early larva. This ectodermal territory of *fz5/8* is complementary to the ectodermal domain of *fz4* in the apical lobe without overlapping with the apical lobe domain of *wnt5* (Fig. 5E,F). *fz5/8* is also expressed in the anterior mesoderm from the early gastrula and in the chaetae sacs of the late larva.

*fz9/10* transcripts are limited to the middle portion of the body throughout development (Fig. 6 and <u>Additional file 9: Fig. S9</u>). In the late blastula, *fz9/10* is initially expressed in the ectoderm posterior to the blastopore, but this domain expands to cover almost the entire ectoderm around the blastopore of the early gastrula; it is only absent from a narrow anterior portion. With gastrulation, *fz9/10* begins to be expressed in the entire mesoderm, as well as in the ectoderm of the apical–mantle boundary, and in the anterior portion of the pedicle lobe ectoderm. Expression in the lateral mantle lobe ectoderm is weaker, and the terminal portion of the pedicle lobe is cleared from *fz9/10* transcripts. Interestingly, the anterior limit of *fz9/10* expression abuts the posterior limit of *fz4* expression in the apical lobe. *fz9/10* expression in the late larva fades, except in the posterior apical lobe ectoderm, and in the anterior and posterior region of the mesoderm.

Taken together, the expression of most Frizzled genes extend over broad but distinct domains along the body. Except for *fz1/2/7*, which is expressed ubiquitously, the ectodermal territories of *fz5/8*, *fz4*, and *fz9/10*, are sequentially arranged from anterior to posterior, respectively, without overlap until the late gastrula stage and the onset of larval morphogenesis.

### Wnt antagonist expression is mostly limited to the anterior end

To obtain a more comprehensive picture of the Wnt signaling landscape in *T. transversa*, we also analyzed the expression of three Wnt antagonist genes: a Secreted Frizzled-Related Protein (*sfrp*), a Dickkopf protein (*dkk*), and a Wnt Inhibitory Factor (*wif*).

sFRP is a soluble protein that antagonizes Wnt activity by directly binding to Wnt ligands or to Frizzled receptors [69]. It has a cysteine-rich domain with high-affinity to Wnt proteins. The sFRP family can be divided into two subfamilies, *sfrp1/2/5* and *sfrp3/4* [69, 70]. In *T. transversa*, we only identified a *sfrp1/2/5* ortholog (<u>Additional file 10: Fig. S10</u>), which is highly expressed throughout development (Fig. 2). The transcripts locate to the animal pole ectoderm in the late blastula and forms a strong anterior ectodermal domain in subsequent stages, in a pattern similar to the expression of *fz5/8* (Fig. 6 and <u>Additional file 11: Fig. S11</u>). *sfrp1/2/5* is also expressed in a narrow domain at the anterior mesoderm throughout development, and in a paired domain in the mantle lobe mesoderm restricted to the early larva stage. In the late larva, the anterior domain becomes limited to dorsal patches on the dorsal ectoderm of the apical lobe.

Dkk is a secreted glycoprotein containing two cysteine-rich domains that antagonizes Wnt signaling by inhibiting *lrp5/6* co-receptors [71, 72]. These proteins are generally divided into two subfamilies, *dkk1/2/4* and *dkk3* [71]. In *T. transversa*, however, we identified a single *dkk* ortholog that groups with a previously unidentified Dkk subfamily, named hereafter *dkk5* (<u>Additional file 12: Fig. S12</u>). Our phylogenetic analysis reveals that non-vertebrate deuterostomes, such as hemichordates and cephalochordates, have orthologs for *dkk1/2/4*, *dkk3*, and *dkk5*, suggesting this was the ancestral Dkk repertoire of bilaterians, and that *dkk1/2/4* and *dkk5* were subsequently lost in protostomes and vertebrates, respectively (<u>Additional file 12: Fig. S12</u>). The expression of *dkk5* in *T. transversa* is upregulated in the late blastula and downregulated in the juvenile (Fig. 2). It localizes to the animal pole ectoderm in the late blastula, and anterior ectoderm in subsequent stages similar to the expression of *sfrp1/2/5*, except that *dkk5* becomes limited to the ventral ectoderm and is not expressed in the mesoderm (Fig. 6 and <u>Additional file 11: Fig. S11</u>).

Wif is another protein that inhibits Wnt activity by direct binding to Wnt proteins [73]. The protein has five EGF repeats and a typical WIF domain which is shared with RYK receptor tyrosine kinases [72, 73]. In *T. transversa*, we identified one *wif* ortholog (<u>Additional file 13: Fig. S13</u>). The expression levels are relatively low and somewhat stable throughout development, with a peak in the late gastrula (Fig. 2). Unlike *sfrp1/2/5* and *dkk5*, *wif* is mainly expressed in mesodermal tissues throughout the analyzed developmental stages; it is also broadly but faintly expressed in the ectoderm until the early larva, and it is not expressed in the endoderm (Fig. 6 and <u>Additional file 11: Fig. S11</u>).

Overall, the expression of the analyzed Wnt antagonist genes is restricted to the anterior portion of the ectoderm (*sfrp1/2/5* and *dkk5*), and to the mesoderm (*wif*), regions which coincide with the absence or limited expression of Wnt ligands.

### Planar cell polarity genes show patched expression during axial elongation

Proper regulation of planar cell polarity (PCP) is crucial to guide morphogenetic processes, such as convergent extension, and to orient the formation of structures during development [74, 75]. We identified several core components of the PCP pathway in *T. transversa*. These include orthologs for *dishevelled* (*dsh*, also known as *dvl*), *diego* (*dgo*, also known as *ankrd6* or *diversin*), *prickle* (*pk*), *flamingo* (*fmi*, also known as *stan* or *celsr*), *strabismus* (*stbm*, also known as *vang* or *vangl*), as well as the downstream transducer *c-jun n-terminal kinase* (*jnk*, also known as *mapk8*). Then, we analyzed their expression between the early and late gastrula stages.

Dsh is a central regulator of the Wnt/beta-catenin, Wnt/PCP, and Wnt/calcium pathways [76]. The protein has three conserved domains (DIX, PDZ, and DEP domains), and two conserved regions before and after the PDZ domain [77]. In *T. transversa*, we identified a single copy of *dsh* (<u>Additional file 14: Fig. S14</u>) which is highly expressed in every developmental stage (<u>Additional file 15: Fig. S15</u>). The expression is stronger in a narrow dorsal domain of the anterior ectoderm and in the anterior portion of the mesoderm (Fig. 7), but *dsh* transcripts are also expressed at lower levels in all embryonic tissues (<u>Additional file 16: Fig. S16</u>).

**Figure 7:**
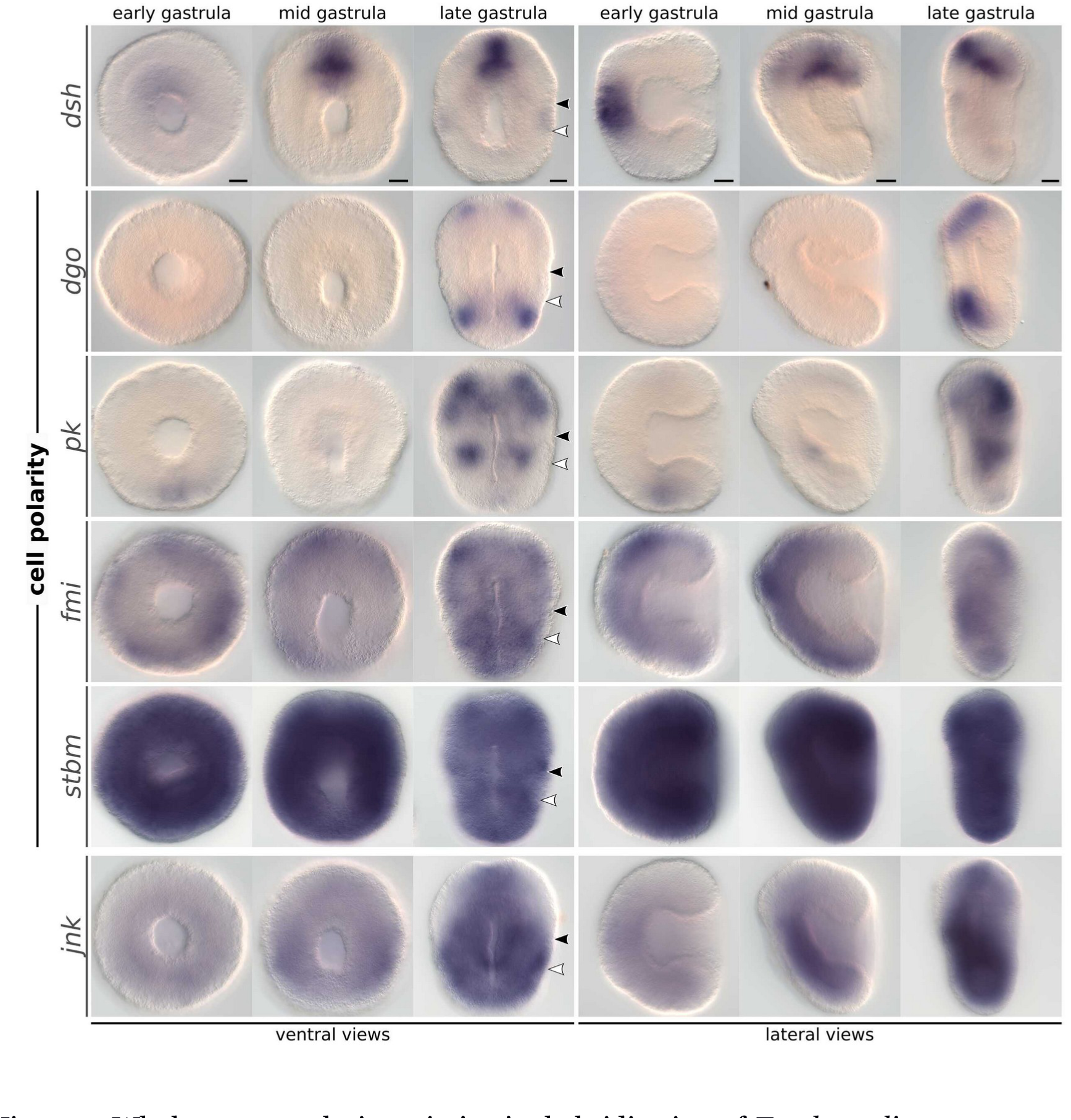
Whole-mount colorimetric *in situ* hybridization of *Terebratalia transversa* Wnt/PCP pathway components. Developmental stages between early gastrula and late gastrula for *dsh*, *dgo*, *pk*, *fmi*, *stbm*, and *jnk*. The panels show representative expression patterns for each sample. The stainings for *dsh* are underdeveloped (see <u>Additional file 16: Fig. S16</u>). The samples are oriented in a blastoporal/ventral view (left) and in a lateral view (right). Black arrowheads indicate the apical–mantle boundary. White arrowheads demarcate the mantle–pedicle boundary. Scale bars = 20µm.

Dgo is a cytoplasmic protein containing 6–8 ankyrin repeat domains that suppresses Wnt/beta-catenin signaling and activates the Wnt/PCP pathway [78, 79]. We found a single *dgo* ortholog in *T. transversa* with six ankyrin repeats (<u>Additional file 17: Fig.</u> <u>S17</u>). *dgo* transcripts are deposited maternally, quickly degrade, and only recover higher levels of expression in the late larva (<u>Additional file 15: Fig. S15</u>). However, we still detect two pairs of dorsal ectodermal domains in the apical and pedicle lobes of the late gastrula (Fig. 7).

Pk is a protein that contains a PET domain and three LIM domains [80] and competes with Dgo for Dsh binding [81]. We identified a single *pk* ortholog in *T. transversa* (<u>Additional file 18: Fig. S18</u>), which is highly expressed throughout development (<u>Additional file 15: Fig. S15</u>). *pk* transcripts are present in a small patch of ectoderm posterior to the blastopore in the early gastrula (Fig. 7). In the mid gastrula, *pk* is upregulated in the mesoderm and forms paired ventral domains within the mantle lobe mesoderm of the late gastrula, when paired ventral domains also appear in the apical lobe ectoderm. Given the high expression levels of *pk* in our RNA-Seq data, we cannot exclude the possibility that it is more broadly expressed than we could detect in our in situ hybridization.

Fmi is a seven-pass transmembrane cadherin that regulates cell polarity [82, 83]. In *T. transversa*, we identified one ortholog of *fmi* (<u>Additional file 19: Fig. S19</u>). In contrast to other polarity genes, it is not expressed maternally; *fmi* expression peaks around the late gastrula (<u>Additional file 15: Fig. S15</u>). *fmi* transcripts are present in most ectodermal tissues but show stronger signal on bilateral patches present in the apical lobe ectoderm of the late gastrula (Fig. 7).

Stbm is a four-pass transmembrane protein that forms a signaling complex with FMI [84, 85]. *Terebratalia transversa stbm* ortholog (<u>Additional file 20: Fig. S20</u>) is initially expressed in high levels, which gradually decay during development (<u>Additional file 15: Fig. S15</u>). Accordingly, *stbm* is ubiquitously expressed in embryonic tissues during gastrulation (Fig. 7).

Jnk is a kinase that regulates epithelial metamorphosis and is a downstream transducer of the PCP pathway [86]. The *jnk* ortholog in *T. transversa* (<u>Additional file 21: Fig. S21</u>) is highly expressed throughout the development (<u>Additional file 15: Fig. S15</u>) and ubiquitously expressed in the late gastrula, except for broad bilateral regions in the apical lobe ectoderm (Fig. 7).

In conclusion, while *fmi*, *stbm*, and *dsh* are expressed ubiquitously, the other cell polarity genes *dgo*, *pk*, and *jnk* are expressed in non-overlapping patches at different regions of the late gastrula.

### Distinct Wnt subregions coincide with larval body subdivisions

Given the importance of specific ligand–receptor contexts for the outcome of Wnt signaling [29, 30], we compiled the data above to describe the combination of Wnt ligands, Frizzled receptors, and antagonist genes being expressed in the different tissues of *T. transversa* embryos throughout ontogeny.

We were able to identify distinct transcriptional subregions, each expressing a unique combination of ligands, receptors, and antagonists, along the brachiopod embryonic axes. At the onset of gastrulation, late blastula embryos exhibit two ectodermal Wnt subregions corresponding to the animal and vegetal tissues (<u>Additional file 22: Fig. S22A</u>). Tissues in the animal pole express Wnt antagonists and all Frizzled genes (except *fz9/10*) but no Wnt genes, while the tissues in the vegetal pole express four ligands (*wnt8* and *wnt16* more broadly) within a *fz1/2/7* and *fz9/10* receptor context. At this stage, the endomesoderm expresses the same ligands as vegetal tissues, except for *wnt4*, but *fz1/2/7* is the only receptor expressed in the invaginating archenteron. From early to mid gastrula, animal and vegetal Wnt subregions subdivide due to changes in the relative position between receptor domains and the upregulation of other ligands. Notably, a subapical Wnt subregion differentiates in the early gastrula, characterized by the expression of *fz4*, *wnt8* and *wnt5*, when *fz5/8* and *fz4* become no longer coexpressed at the animal pole (<u>Additional file 22: Fig. S22B</u>). Likewise, two Wnt subregions emerge in the mid gastrula from the initial vegetal landscape when *fz9/10* expression becomes subterminal—one midbody, continuing to express *fz1/2/7*, *fz9/10*, *wnt7*, and *wnt16*, and one posterior, expressing only *fz1/2/7* and several Wnt genes (<u>Additional file 22: Fig. S22C</u>). Finally, at the late gastrula stage, *fz9/10* is cleared from a portion of the midbody and *fz1/2/7* is cleared from the posterior end, giving rise to an additional midbody subregion expressing *fz1/2/7*, *wnt5* and *wnt16*, and to another subregion at the posterior end expressing only Wnt but no Frizzled genes (Fig. 8A). Therefore, from late blastula to the elongated late gastrula, we observe a progressive differentiation of ectodermal Wnt subregions, from the initial animal and vegetal ones, to the six distinct subregions along the anteroposterior axis of the embryo (Fig. 8A, Table 2).

**Figure 8:**
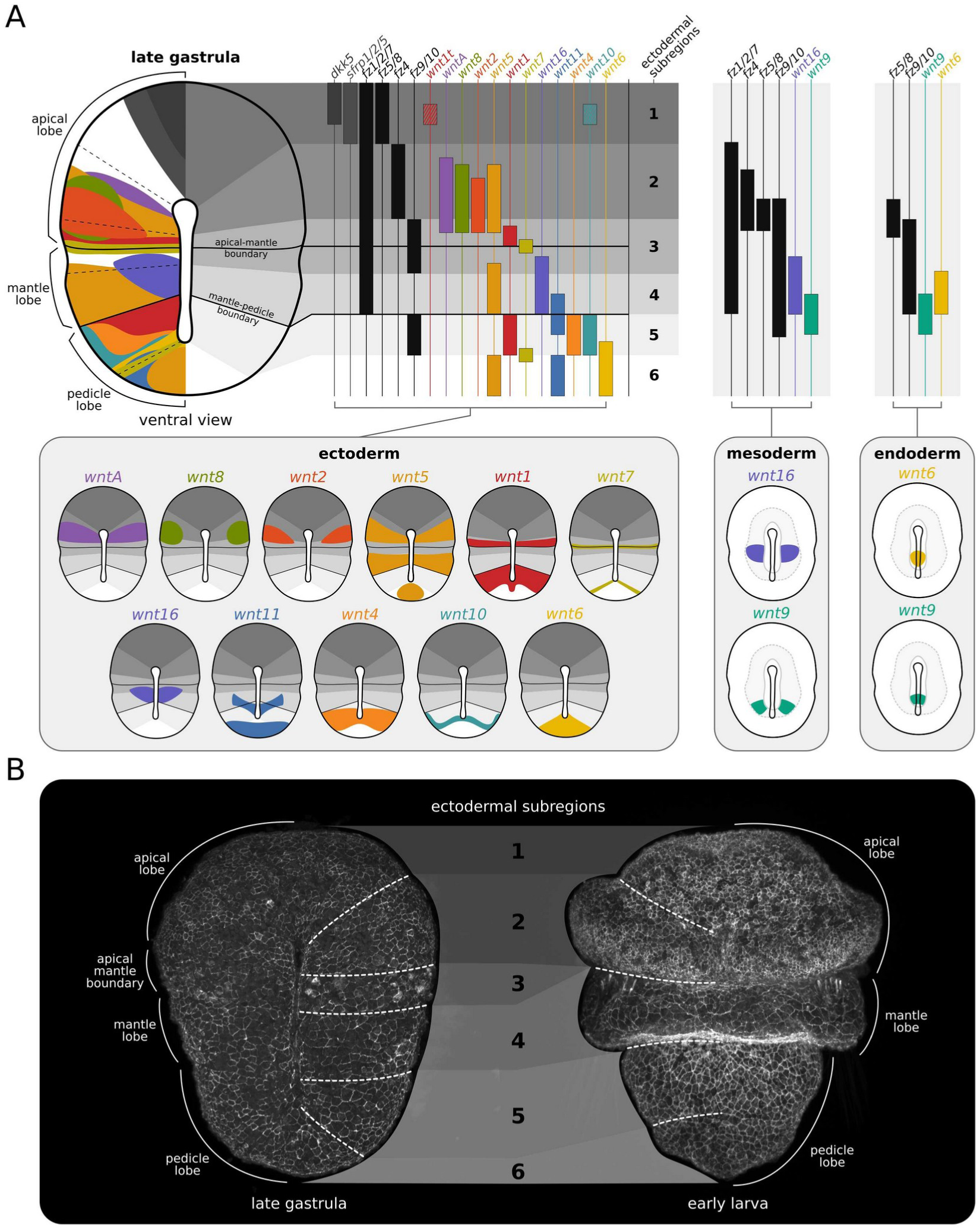
Summary of *Terebratalia transversa* Wnt signaling landscape during axial elongation. (A) Late gastrula in blastoporal/ventral view. Schematic drawings of Wnt genes colored by subfamilies, Frizzled genes by lighter shades of gray, and antagonists by darker shades of gray. The spatial localization of expression domains is superimposed on the embryo (left) and projected to highlight the individualized Wnt genes within the different transcriptional subregions grouped by germ layer (right). The gray boxes show the pattern of individual genes mapped to the embryo for clearer visualization of overlapping domains. (B) Late gastrula and early larva embryos showing the correspondence between Wnt subregions (dashed lines) and the morphological boundaries of the larva. The samples were stained for F-actin (cell membranes) to highlight the cell shape differences between body regions.

**Table 2:** *Wnt signaling transcriptional subregions in the brachiopod Terebratalia transversa*.

| Subregion | Position | Antagonists | Receptors | Ligands |
| --- | --- | --- | --- | --- |
| 1 | Anterior portion of apical lobe | *dkk*, *sfrp1/2/5* | *fz1/2/7*, *fz5/8* | *wnt1t* |
| 2 | Posterior portion of apical lobe | - | *fz1/2/7*, *fz4* | *wntA*, *wnt8*, *wnt2*, *wnt5* |
| 3 | Boundary between apical and mantle lobes | - | *fz1/2/7*, *fz9/10* | *wnt1*, *wnt7*, *wnt16* |
| 4 | Mantle lobe | - | *fz1/2/7* | *wnt5*, *wnt16* |
| 5 | Anterior portion of pedicle lobe | - | *fz9/10* | *wnt1*, *wnt9*, *wnt11*, *wnt4*, *wnt10*, *wnt7* |
| 6 | Posterior portion of pedicle lobe | - | - | *wnt6*, *wnt11*, *wnt5* |

Remarkably, the Wnt subregions established in the late gastrula stage coincide with the morphological subdivisions of the larval body (Fig. 8B). The anteriormost subregion expressing antagonist genes and *fz5/8* but no Wnt ligands (except *wnt1t*), gives rise to the larval apical organ and neuropile at the narrower portion of the apical lobe [87]. In turn, the adjacent subapical subregion expressing Wnt genes and *fz4* but no antagonists, undergoes intense cell proliferation [63] and gives rise to the wider portion of the larval apical lobe and adult lophophore, the crown of tentacles that brachiopods use for filter-feeding (Fig. 8). The subregion posterior to the apical lobe, which expresses *fz1/2/7*, *fz9/10*, *wnt1*, and *wnt7*, gives rise to a deep epithelial invagination that demarcates the apical–mantle boundary [64]. The abutting subregion expressing solely *fz1/2/7*, *wnt5*, and *wnt16*, gives rise to a prominent, skirt-like epithelial outgrowth that forms the larval mantle lobe, a structure that is then reversed anteriorly during metamorphosis [53] (Fig. 8). This reversible mantle lobe is considered an evolutionary innovation of lecithotrophic brachiopod larvae [88]. Tissues posterior to the mantle lobe, which express *fz9/10*, *wnt1*, *wnt4*, and *wnt10*, become the posterior end of the adult body, while the posteriormost subregion expressing *wnt5*, *wnt6*, and *wnt11*, but no Frizzled genes, gives rise to the posterior end of the larval pedicle lobe which becomes the structure that attaches adult brachiopods to the substrate [54] (Fig. 8). Overall, a combinatorial analysis of the expression of Wnt signaling components in *T. transversa* reveals that the distinct transcriptional subregions established during gastrulation correlate with the distinct morphogenetic outcomes along the anteroposterior axis of larval brachiopods.

## Discussion

Our work characterizes the expression of Wnt signaling components during the embryonic development of *T. transversa*, a brachiopod species that has largely retained the ancestral repertoire of Wnt genes. We find that Wnt genes are upregulated at the onset of gastrulation and during axial elongation, and that the combinatorial expression of ligands, receptors, and antagonists, forms distinct transcriptional subregions along the primary body axis. The boundaries between these subregions coincide with the morphological subdivisions of the larval body, suggesting that a combinatorial Wnt signaling landscape—or *Wnt code*—may play a role in the regionalization of the brachiopod anteroposterior body axis.

We observe, for example, that the expression of Wnt inhibitors in *T. transversa* correlates with the anteriormost fates of the apical lobe in the brachiopod larva (Fig. 8). In addition to *sfrp* and *dkk5*, the anterior subregion of the brachiopod also expresses *fz5/8*, a gene known to activate sFRP and Dkk antagonists in sea urchins [89, 90], and *six3/6* [58], a transcription factor that antagonizes Wnt/beta-catenin signaling in different animals [91–94]—but almost no Wnt genes. This could indicate that Wnt/ beta-catenin signaling is being inhibited at the anteriormost brachiopod subregion, a hypothesis consistent with the observation that early over-activation of this pathway in *T. transversa* dramatically reduces the expression of anterior markers and results in the complete loss of anterior structures (apical and mantle lobes) (see Supplementary Figure S9 in [64], and Figure 4d and Supplementary Figure 6b in [63]). Interestingly, the adjacent subregion in the brachiopod expresses *wnt8* and other ligands posteriorly (<u>Additional file 22: Fig. S22</u>), a pattern comparable to the expression of Wnt components in the anterior neuroectoderm of sea urchin embryos [95–97]. In echinoderms, this pattern emerges from a negative feedback loop between the anterior Wnt inhibitors (activated by *fz5/8*), and the posterior *wnt8*-mediated Wnt/betacatenin signaling, which helps consolidate the anterior and posterior neuroectoderm identities [89, 90]. Whether a comparable regulatory logic could also be contributing to the patterning of the brachiopod neuroectoderm and apical lobe subdivision remains a speculation at this point, and will need to be experimentally tested in future studies.

Our study also reveals a correlation between *wnt5* and the formation of the mantle lobe in *T. transversa*. The Wnt subregion that gives rise to the mantle lobe is characterized by the expression of *wnt5* and *fz1/2/7* and emerges only in the late gastrula with the clearance of *fz9/10* transcripts from embryo’s midbody (Fig. 8)—*wnt16* is also expressed but limited to the midline around the closing blastopore. During larval morphogenesis, *wnt5* remains expressed at the growing edges of the elongating mantle lobe (Fig. <u>Additional file 3: Fig. S3</u>). In other animals, *wnt5* regulates convergent extension movements during tissue morphogenesis via the Wnt/PCP pathway [3, 16, 17, 98], and is commonly expressed in tissue outgrowths such as the vertebrate tail and limb buds [23, 99, 100]. This contextual similarity of *wnt5* expression suggests that convergent extension could be a possible mechanism for the elongation of the mantle lobe in brachiopod larvae. One piece of evidence consistent with this hypothesis is that a late over-activation of the Wnt/beta-catenin pathway in *T. transversa* inhibits the elongation of the mantle lobe (see Supplementary Figure S8 in [64]). However, whether *wnt5* or the Wnt/PCP pathway plays a role in this process, and whether the specific receptor context in the brachiopod mantle lobe is important to control the signaling output, remains to be determined by targeted functional approaches.

Altogether, these observations suggest the possibility that the different transcriptional subregions could contribute to the regional specification and morphogenetic control of tissues along the anteroposterior axis of brachiopods. It is important to note, however, that these combinatorial patterns are solely based on coexpression data, which does not guarantee actual signaling, and that long-range interactions might still occur due to the secreted nature of Wnt proteins and their highly promis-cuous ligand–receptor binding. Nevertheless, our brachiopod data hints at the idea that changes in these Wnt subregions may have been important for developmental innovations in the primary body axis of animals.

A broad phylogenetic survey comparing our brachiopod data to other animals suggests that the combinatorial expression of Wnt signaling components along the body emerged with the Cnidaria–Bilateria expansion Wnt and Frizzled gene families (Fig. 9A). Ctenophores have only four Wnt, two Frizzled, and one sFRP and their expression is not staggered along the embryonic ectoderm [101]. Sponges exhibit Wnt territories along the larval body axis [102–104], but the expression data is scarce and some species underwent large Wnt expansions which have no clear orthologs with the bilaterian Wnt genes [103, 104]. While the early evolution of Wnt genes remains poorly understood, the Wnt repertoire between cnidarians and bilaterians is well conserved with clear orthologs, enabling a more informative comparison about the evolution of the Wnt signaling landscape (Fig. 9A).

**Figure 9:**
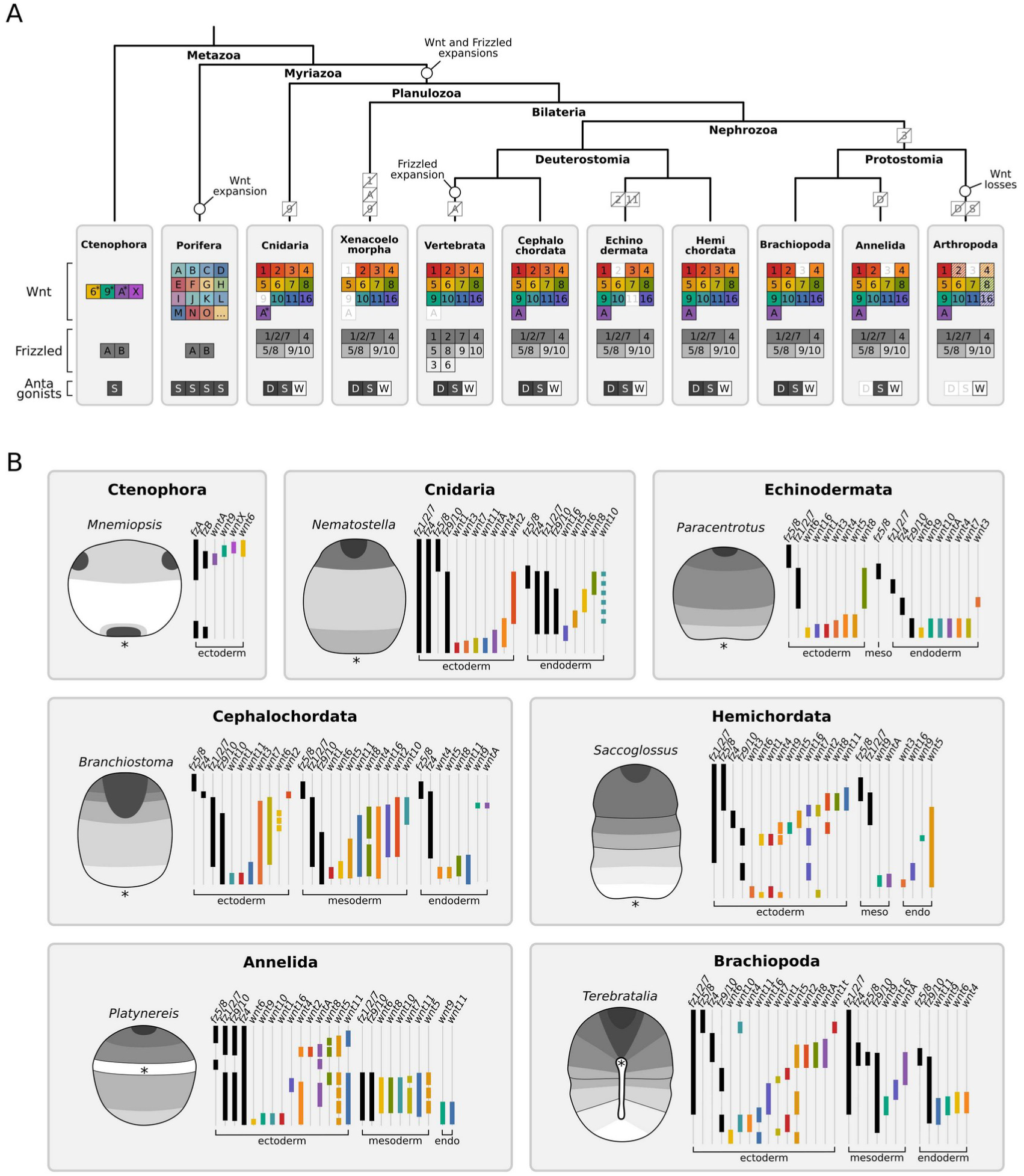
Comparative overview of the Wnt signaling landscape in animals. (A) Phylogenetic tree showing the repertoire of Wnt ligands, Frizzled receptors, and Wnt antagonists in different animal groups, and their presumed gene expansions and gene losses during evolution. Ctenophores have few Wnt genes while poriferans, due to lineage-specific expansions, have several Wnt genes; their orthology to bilaterian Wnt genes, however, remains uncertain. For this reason, it is assumed that Wnt and Frizzled genes expanded to 13 and 4 subfamilies, respectively, in the last common an-cestor of cnidarians and bilaterians. Alternatively, ancestral orthologs of Wnt subfamilies may have been lost or significantly modified in ctenophores and sponges. The main losses in bilaterians were *wntA* in vertebrates, *wnt2* and *wnt11* in echinoderms, *wnt3* in protostomes, and multiple Wnt genes in arthropods—but, generally, they show a well-conserved Wnt repertoire. Asterisks indicate Wnt genes with uncertain orthology. The tree topology is based on [127, 128]. (B) The schematic drawings illustrating the expression of Wnt signaling components at the late gastrula stage of different metazoan species. The receptor (lighter shades of gray) and antagonist (darker shades of gray) subregions are superimposed on the embryo (left). The region of expression of individual Frizzled genes (black lines) and Wnt genes (colored lines) is shown for each species depicted by germ layer. Embryos are oriented with aboral/ anterior end up and oral/posterior end bottom. Asterisk indicates the blastopore position. The gene complement, orthology assignment, and expression patterns were compiled from previous works on Ctenophora [101, 129], Porifera [102–104], Cnidaria [24, 25, 130, 131], Xenacoelomorpha (based on *Xenoturbella* transcriptome [132] as the Wnt genes in acoels are highly derived [133]), Vertebrata [1], Cephalo-chordata [23, 105, 134–140], Echinodermata [68, 89, 90, 95–97, 141–143], Hemi-chordata [41], Brachiopoda (based on [64] and this study), Annelida [27, 40, 47, 48, 144], and Arthropoda [44].

Despite the great morphological differences, most both cnidarian and bilaterian embryos exhibit a common spatial organization of their Wnt landscape with antagonist genes expressed anteriorly, Wnt genes expressed predominantly posteriorly, and Frizzled genes expressed more broadly along the body (Fig. 9B). Frizzled expression is especially well conserved across groups. While most animals provide maternal *fz1/2/7* and *fz5/8* transcripts in the egg (Table 3), embryos of every group investigated so far express *fz5/8* at the aboral/anterior ectoderm, often coexpressed with *sfrp* and rarely with Wnt ligands (Fig. 9B). Expression of *fz5/8* in the anterior mesoderm is also common. *fz9/10* expression is usually complementary to *fz5/8*, localizing to the midbody/trunk portion of the body, and *fz1/2/7* is normally expressed broadly, even though the anterior and posterior domain limits can vary (Fig. 9B). A closer comparison between *T. transversa* and the hemichordate *Saccoglossus kowalevskii* reveals that the spatial organization of their Frizzled domains is strikingly similar (<u>Additional file 23: Fig. S23A</u>). The exception to this conservation is *fz4*, which is subapical in brachiopods and hemichordates, but shows a more variable expression in other groups. Overall, these similarities indicate that cnidarians and bilaterians share an ancient axial organization of Frizzled domains that remained conserved during evolution.

**Table 3:** *Maternal load of Wnt and Frizzled transcripts in metazoan eggs*.

| Group | Species | Frizzled genes | Wnt genes |
| --- | --- | --- | --- |
| Brachiopoda | *Terebratalia transversa* | *fz1/2/7, fz5/8* and *fz4* (this study) | Absent (this study) |
| Brachiopoda | *Lingula anatina* | *fz1/2/7, fz4* and *fz9/10* [145, 146] | *wntA* and *wnt8* [145, 146] |
| Annelida | *Platynereis dumerilii* | *fz1/2/7* [144] | Absent [40] |
| Priapulida | *Priapulus caudatus* | - | *wnt2, wnt4, wnt5* and *wnt8* [147] |
| Hemichordata | *Saccoglossus kowalevskii* | *fz1/2/7* and *fz5/8* [41] | *wnt4* and *wnt9* [41] |
| Echinodermata | *Paracentrotus lividus* | *fz1/2/7* and *fz5/8* [96] | *wnt7* [96] |
| Echinodermata | *Strongylocentrotus purpuratus* | *fz1/2/7* [141] | *wnt6, wnt7* and *wnt16* [148] |
| Vertebrata | *Xenopus laevis* | - | *wnt5, wnt8* and *wnt11* [149] |
| Vertebrata | *Danio rerio* | - | *wnt8* [150] |
| Cnidaria | *Nematostella vectensis* | *fz1/2/7* and *fz5/8* [131] | - |
| Cnidaria | *Clytia hemisphaerica* | *fz1/2/7* and *fz9/10* [151] | *wnt3* [152] |

In contrast, the expression of Wnt genes is more variable. The maternal load of Wnt transcripts is diverse and can even differ within a group (Table 3), while the embryonic expression shows great variability across the body axis and germ layers between groups (Fig. 9B). *wnt2* is the least variable gene. Except for echinoderms which lack the gene, *wnt2* is the only Wnt to be solely expressed in a single body region and germ layer across groups—the anterior ectoderm. *wnt1* has the most consistent expression domain at the posterior ectoderm, but it is also expressed at the anterior ectoderm (Fig. 9B and <u>Additional file 23: Fig. S23A</u>). The other frequent ectodermal patterns are *wnt5* and *wnt8* in the anterior, and *wnt4*, *wnt3*, *wnt6*, and *wnt11* at the posterior end. However, these genes are also expressed in other positions and germ layers. The most diverse patterns in this sense are from the genes *wnt11*, *wnt9*, and *wnt5* (Fig. 9B). *wnt9* and *wnt11*, for example, are expressed at entirely different anterior and posterior receptor contexts between brachiopod and hemichordate embryos (<u>Additional file 23: Fig. S23A</u>). In fact, based on our expression data, even the consistent genes *wnt1* and *wnt2* are expressed in different receptor contexts between groups. A more detailed comparison between the brachiopod and hemichordate landscapes reveals only a few relatively conserved Wnt–Frizzled combinations (<u>Additional file 23: Fig. S23B</u>). Altogether, this comparative analysis suggests that changes in Wnt expression, like the addition or re-deployment of domains to different tissues or receptor contexts, may have occurred frequently during evolution.

This variable nature of Wnt expression might be an important factor influencing axial evolution. Evolutionary changes in the ligand–receptor context could effectively alter cell fates and morphogenetic events, generating tissue and shape diversity and providing a basis for developmental innovations along the body [3, 25, 26]. Such Wnt function shuffling is indeed associated with novelties in the chordate lineage [105]. Investigating the combinatorial landscape of Wnt signaling components across the phylogeny will be crucial to uncovering the role of Wnt shuffling as a potential mechanism contributing to the diversification of the metazoan body axis.

## Conclusions

Our data reveals a correlation between distinct Wnt transcriptional subregions and the morphological subdivisions in the larval body of a brachiopod. We hypothesize that the underlying combinatorial landscape may play a role in the patterning and morphogenesis of the different regions along the anteroposterior axis, and that changes in this landscape could be associated with the evolution of the reversible mantle lobe, a morphogenetic innovation of brachiopod larvae. As Wnt developmental expression is variable across animal groups, we propose that evolutionary changes in ligand–receptor context may have been important to axial evolution in animals.

## Methods

### Sample collection

*T. transversa* (Sowerby, 1846) adult specimens were collected by dredging the rocky seabeds around Friday Harbor, San Juan Islands, USA. We kept in a seawater table with running seawater at the Friday Harbor Laboratories (University of Washington). To obtain embryos, we dissected the gonads of ripe individuals and fertilized the gametes *in vitro* as previously described [56, 106]. We cultured the embryos in E-ware glass bowls (i.e., glassware never exposed to chemicals) with filtered seawater and temperature around 7.6 °C. Water in culturing bowls was exchanged daily. Using a glass pipette, we collected samples for RNA sequencing and for *in situ* hybridization at representative developmental stages (Table 1). For the RNA-Seq samples, we collected two biological replicates, each containing the eggs of a single female fertilized with mixed sperm from three different males. We preserved the embryos directly in RNAlater at room temperature. For the *in situ* hybridization samples, we fixed the embryos for 1h in 4% paraformaldehyde at room temperature, washed thoroughly in 1x PBS with 0.1% Tween-20, and stored them in 100% methanol at -20°C.

### RNA sequencing and analyses

We extracted the total RNA using Trizol. Library preparation and sequencing was performed at the EMBL Genomic Core Facilities (GENECORE). The samples were randomized and multiplexed on four lanes of a Illumina HighSeq 2000 system, and sequenced to an average of 24±5 million 50bp of unstranded single-end reads. To quantify transcript abundances, we used Kallisto v0.46.0 [107] to pseudoalign the reads to a reference transcriptome of *T. transversa*. This reference transcriptome was originally assembled with Trinity [108] from a deeply sequenced, unstranded paired-end RNA-Seq dataset of mixed developmental stages [109]. Next, we imported the estimated counts from Kallisto to DESeq2 [110] to estimate the library size factors and data dispersion, homogenize the variance across expression ranks, and apply a variance-stabilizing transformation to the data before the expression analyses. To visualize the normalized expression data, we generated heatmaps using pheatmap [111] and ggplot2 [112] in R [113] running in RStudio Desktop [114].

Due to the unstranded nature of our sequencing data, we analyzed the coverage of mapped reads to identify potential biases in the quantification of expression levels. For that, we mapped the RNA-Seq reads to the transcripts of Wnt signaling components of *T. transversa* using Salmon v1.10.1 [115], and then created read coverage plots for each gene using the ggcoverage package [116]. While the majority of genes show uniform read coverage profiles, we identified two cases of uneven coverage that significantly overestimates the expression levels of *wnt4* and *wntA* in early developmental stages (<u>Additional file 24: Fig. S24</u>). In these samples, reads predominantly mapped to the 3’ region of the transcript, while the Wnt coding sequence region had a low mapping rate. This pattern could be explained by the presence of an isoform lacking the Wnt domain, or by the expression of another gene in the opposite strand at the same locus, potentially assembled in the same contig due to the unstranded reads. Since both *wnt4* and *wntA* transcripts have long open reading frames in the antisense direction, and these transcripts fully map to a single scaffold in a draft genome assembly of *T. transversa*, the latter hypothesis is more likely. This suggests that the high expression values result from the contiguous antisense gene rather than *wnt4* or *wntA*.

The code and pipeline for the RNA-Seq analysis and the mapping and coverage files are available in the paper’s repository [117].

### Gene orthology

We searched for Wnt signaling orthologs in *T. transversa* by querying known Wnt genes against the available transcriptome using BLAST+ [118]. To resolve the orthology of the obtained putative orthologs, we aligned the protein sequences of *T. transversa* with well-annotated proteins from other metazoans using MAFFT 7.310 [84], removed non-informative sections using GBlocks 0.91b [119], and inspected the multiple sequence alignment using UGENE [120]. Using the blocked alignments as input, we ran a maximum likelihood phylogenetic analysis using the automatic model recognition and rapid bootstrap options of RAxML 8.2.12 [121], and rendered the resulting trees using the Interactive Tree Of Life web application [122]. The gene orthology pipeline is available in the paper’s repository [117].

### Cloning and *in situ* hybridization

We synthesized cDNA from a total RNA extraction of mixed developmental stages of *T. transversa* using the SMARTer RACE cDNA Amplification kit (Clontech). For each transcript, designed gene-specific primer pairs within the coding sequence using Primer3 [123] to obtain product sizes between 800 and 1200bp (<u>Additional file 25: Table S1</u>). We then cloned each fragment into a pGEM-T Easy Vector, amplified the antisense sequences using T7 or SP6 polymerase, and synthesized DIG-labeled riboprobes using the MEGAscript kit (Ambion). Finally, to visualize gene expression, we followed the established protocols in *T. transversa* for single colorimetric *in situ* hybridization [58, 124], and double fluorescent *in situ* hybridization [63, 64]. Cloning details are available in the paper’s repository [117].

### Microscopy and image processing

We mounted the embryos between a glass slide and a coverslip, supported by clay feet, in 70% glycerol in PBS, and imaged the representative expression patterns for each sample. For colorimetric *in situ* data, we acquired differential interference contrast (DIC or Nomarski) images using a Zeiss AxioCam HRc attached to a Zeiss Axioscope A1. For fluorescent samples, we scanned volumetric stacks in a Leica TCS SP5 confocal microscope and generated maximum intensity projections using Fiji [125]. We adjusted intensity levels to improve contrast—without clipping signal from high or low ranges—using Fiji or GIMP, and assembled the illustrations and figure plates using Inkscape. Images at the original resolution are available in the paper’s repository [117].

## Declarations

### Acknowledgments

We thank the Friday Harbor Laboratories boat crew for collecting the brachiopods, Yale Passamaneck for the help with spawnings, Katrine Worsaae and Yvonne Müller for initial gene cloning, Juliana Roscito for the help with coverage analysis, and members of the Hejnol Lab for the productive discussions. We also thank Grigory Genikhovich and two anonymous reviewers for their insightful comments and constructive feedback. BCV thanks Pavel Tomančák for the generous support during the preparation of this work.

### Consent for publication

Not applicable.

### Funding

The study was funded by the Michael Sars Centre core budget and by The European Research Council Community’s Framework Program Horizon 2020 (2014–2020) ERC grant agreement 648861 to AH. The animal collections were supported by a

Meltzer Research Fund. BCV was funded by an EMBO Long-Term Fellowship (ALTF 74-2018) during the writing of this manuscript.

### Availability of data and materials

All data generated or analyzed during this study are included in this published article, its supplementary information files, and publicly available repositories.

- **Paper’s repository:** Zenodo https://doi.org/10.5281/zenodo.8312022. Citable repository containing the source code, data files, analysis pipelines, figure plates, manuscript text, original *in situ* images, and high-resolution figures [117].
- **Stage-specific RNA-Seq:** NCBI https://identifiers.org/bioproject:PRJNA1103701. Raw reads deposited as BioProject PRJNA1103701 and SRA experiments SRX24343897–SRX24343924 [126].
- **Gene sequences:** NCBI. mRNA sequences deposited with the accession numbers KT253961, PP860497–PP860521.

### Authors’ contributions

AH, JMMD, and BCV designed the study and collected the samples. BCV, JMMD, and AB performed gene cloning, *in situ* hybridization, and imaging. BCV performed the phylogenetic and transcriptomic analyses. BCV analyzed the data, prepared the figures, and wrote the manuscript. AH and JMMD revised and contributed to the text. All authors read and approved the final manuscript.

### Competing interests

The authors declare that they have no competing interests.

### Ethics approval and consent to participate

Not applicable.

## Abbreviations

DGO: diego
DKK: dickkopf
DSH: dishevelled
FMI: flamingo
FZ: frizzled
JNK: c-jun n-terminal kinase
PCP: planar cell polarity
PK: prickle
SFRP: secreted frizzled-related protein
STBM: strabismus
WIF: wnt inhibitory factor

## Supplementary information

**Additional file 1: Fig. S1:**
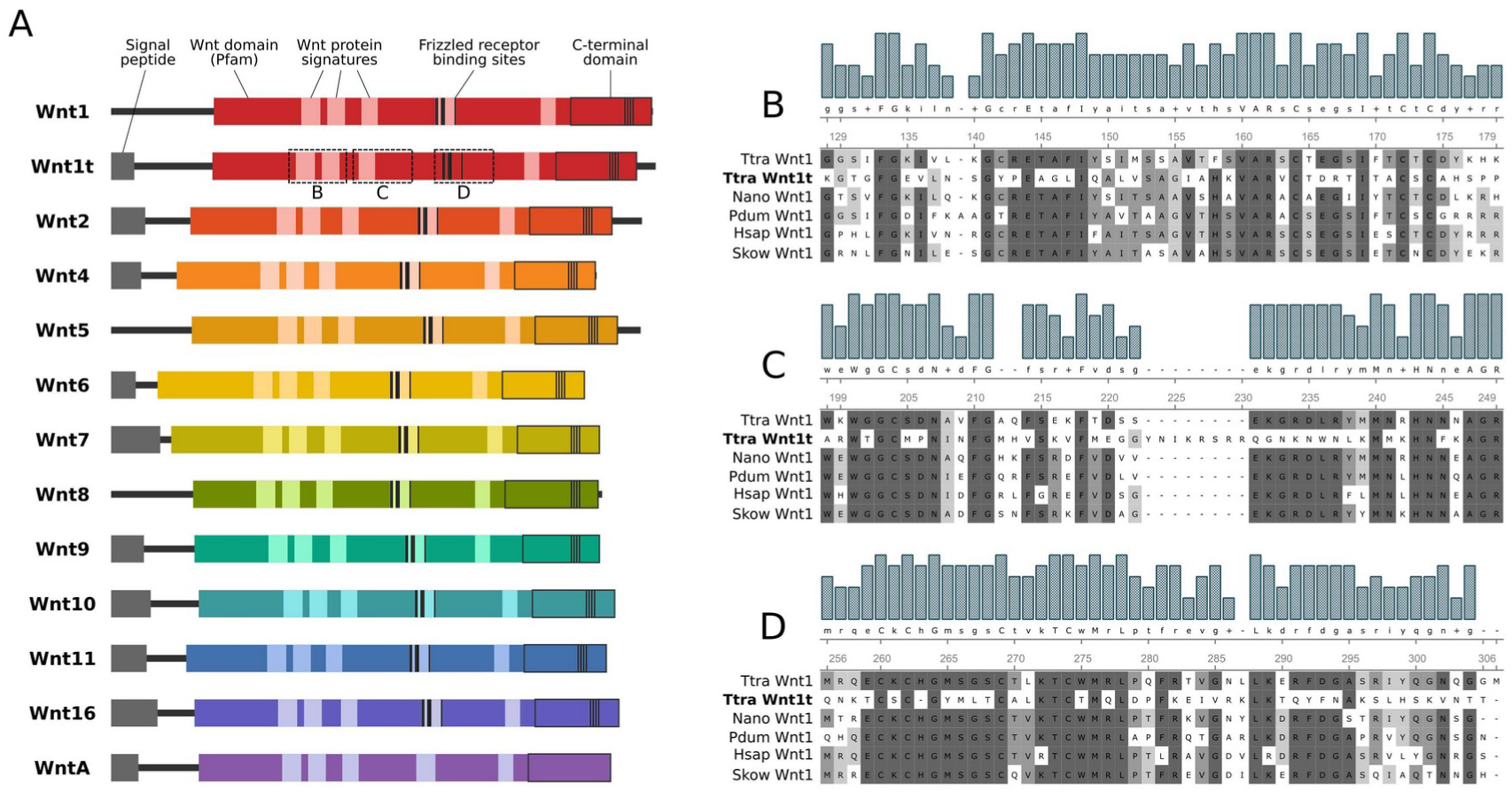
[PDF] Domain architecture of *Terebratalia transversa* Wnt proteins. (A) Schematic drawings showing signal peptide regions, Wnt protein signatures, Frizzled-receptor binding sites, and C-terminal Wnt domain based on InterProScan annotations. All *T. transversa* have a similar overall architecture. (B–C) Multiple sequence alignment of Wnt1 proteins, showing the highly divergent sequence of *T. transversa* Wnt1t in three Wnt protein signature regions. The alignment contains Wnt1 orthologs of *T. transversa* (Ttra), *Novocrania anomala* (Nano), *Platynereis dumerilii* (Pdum), *Homo sapiens* (Hsap), and *Saccoglossus kowalevskii* (Skow).

**Additional file 2: Fig. S2:**
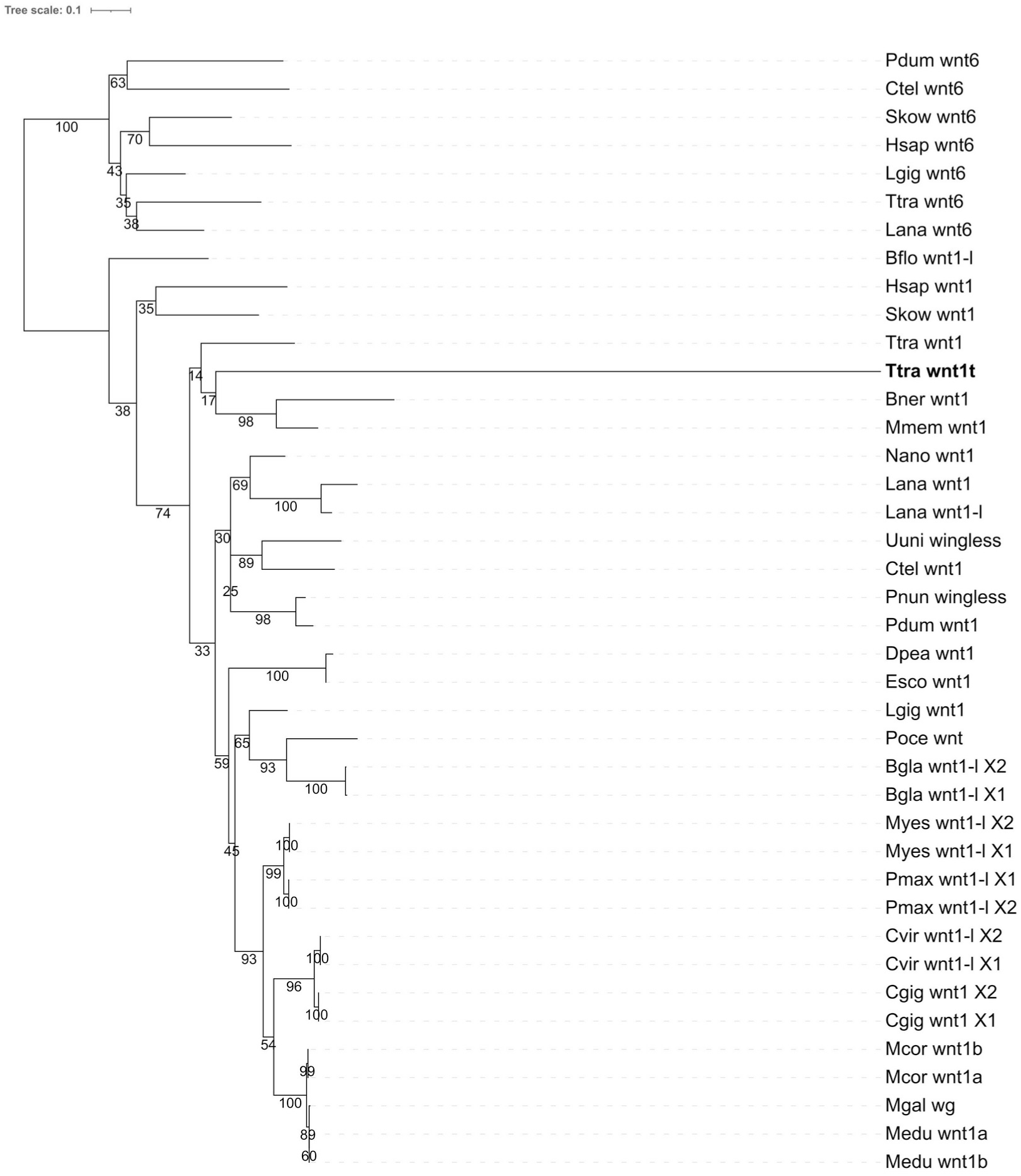
[PDF] Phylogenetic analysis of *Terebratalia transversa* Wnt1 proteins. Best-scoring tree of a maximum likelihood phylogenetic analysis using amino acid sequences of genes from the *wnt1* subfamily with *wnt6* as an outgroup. Branch lengths are proportional to the amount of sequence change, and the numbers show the support values of individual branches. Both *Terebratalia transversa* (Ttra) and *Lingula anatina* (Lana), a rhynchonelliform and a linguliform brachiopod, respectively, have two copies of *wnt1*. If this was an ancient duplication event at the base of Brachiopoda, we would expect the orthologous *wnt1* paralogs from different species to cluster together (i.e., Ttra *wnt1* with Lana *wnt1*). Instead, the tree reveals that the paralog copies of each species cluster together, suggesting that the duplication of *wnt1* occurred independently in *T. transversa* and *L. anatina*. *T. transversa wnt1t* also shows a longer branch length indicating rapid evolution. Taxon sampling was focused in spiralians. The other species are *Biomphalaria glabrata* (Bgla), *Branchiostoma floridae* (Bflo), *Bugula neritina* (Bner), *Capitella teleta* (Ctel), *Crassostrea virginica* (Cvir), *Doryteuthis pealeii* (Dpea), *Euprymna scolopes* (Esco), *Homo sapiens* (Hsap), *Lingula anatina* (Lana), *Lottia gigantea* (Lgig), *Membranipora membranacea* (Mmem), *Mizuhopecten yessoensis* (Myes), *Mytilus coruscus* (Mcor), *Mytilus edulis* (Medu), *Mytilus galloprovincialis* (Mgal), *Pecten maximus* (Pmax), *Perinereis nuntia* (Pnun), *Plakobranchus ocellatus* (Poce), *Platynereis dumerilii* (Pdum), *Saccoglossus kowalevskii* (Skow), and *Urechis unicinctus* (Uuni).

**Additional file 3: Fig. S3:**
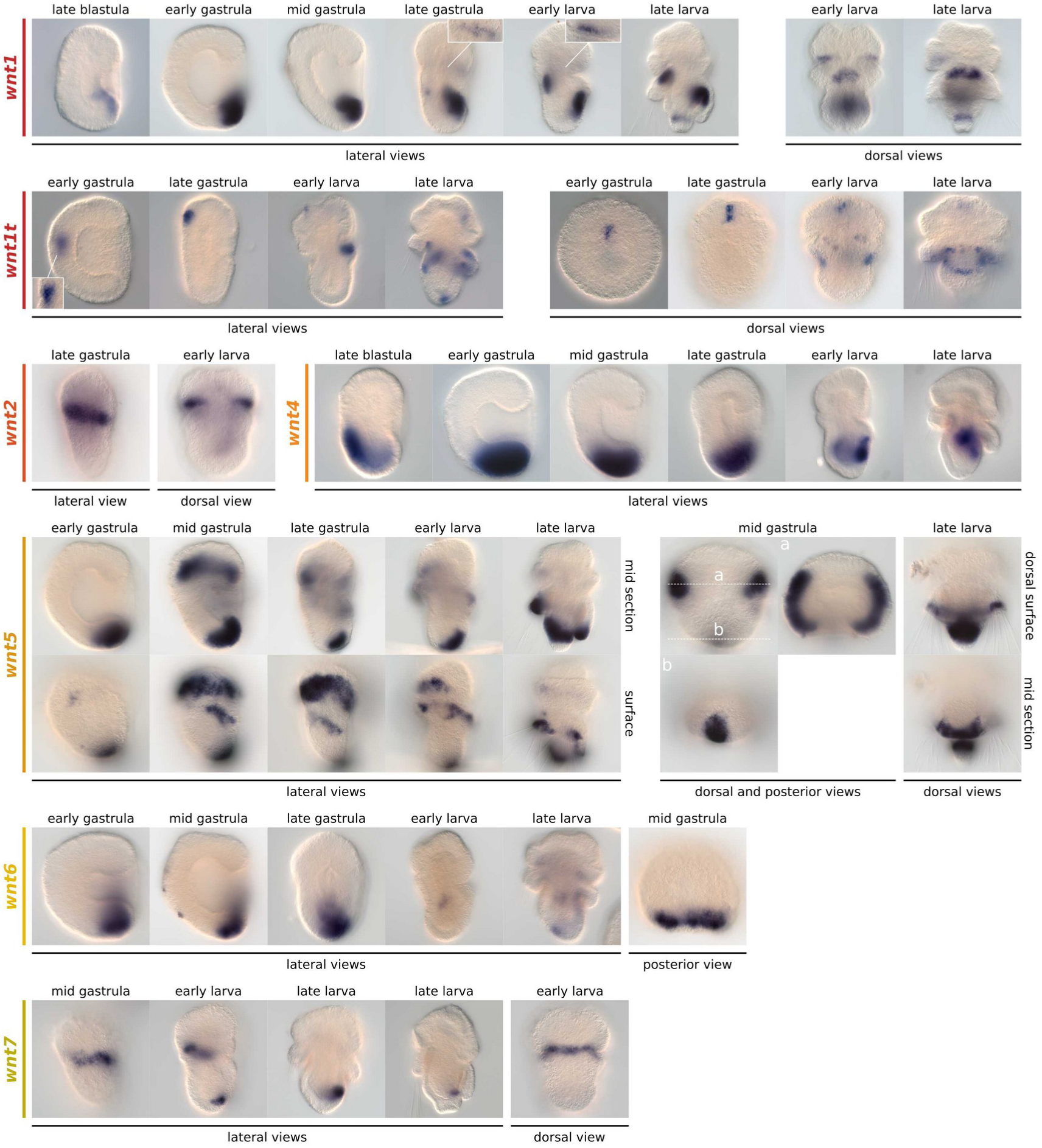
[PNG] Whole-mount colorimetric *in situ* hybridization of *wnt1*, *wnt1t*, *wnt2*, *wnt4*, *wnt5*, *wnt6*, and *wnt7* in *Terebratalia transversa*. Additional views of Wnt expression between late blastula and late larva. Dashed lines indicate the position of the optical section shown in adjacent panels. The panels show representative expression patterns for each sample.

**Additional file 4: Fig. S4:**
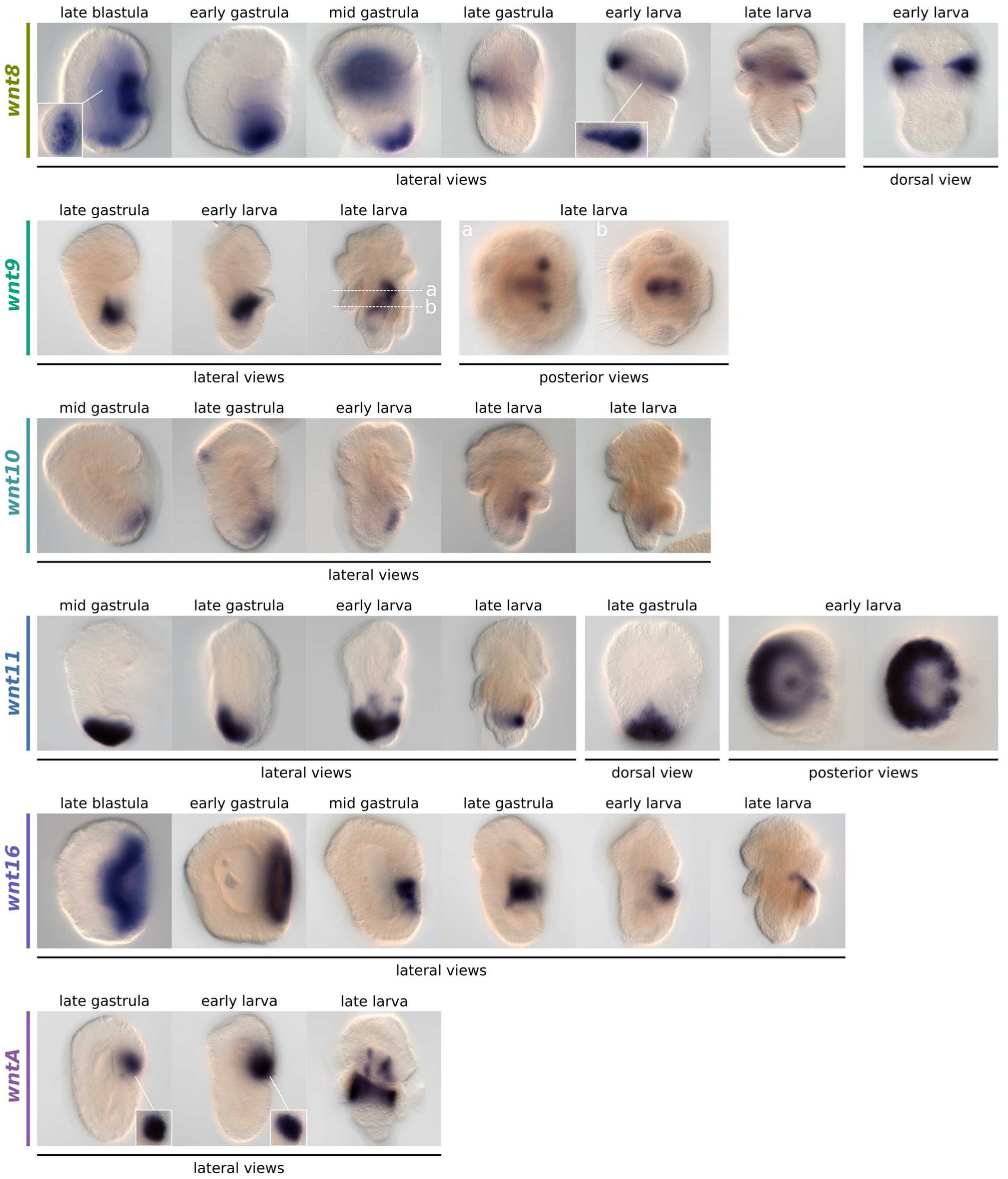
[PNG] Whole-mount colorimetric *in situ* hybridization of *wnt8*, *wnt9*, *wnt10*, *wnt11*, *wnt16*, and *wntA* in *Terebratalia transversa*. Additional views of Wnt expression between late blastula and late larva. Dashed lines indicate the position of the optical section shown in adjacent panels. The panels show representative expression patterns for each sample.

**Additional file 5: Fig. S5:**
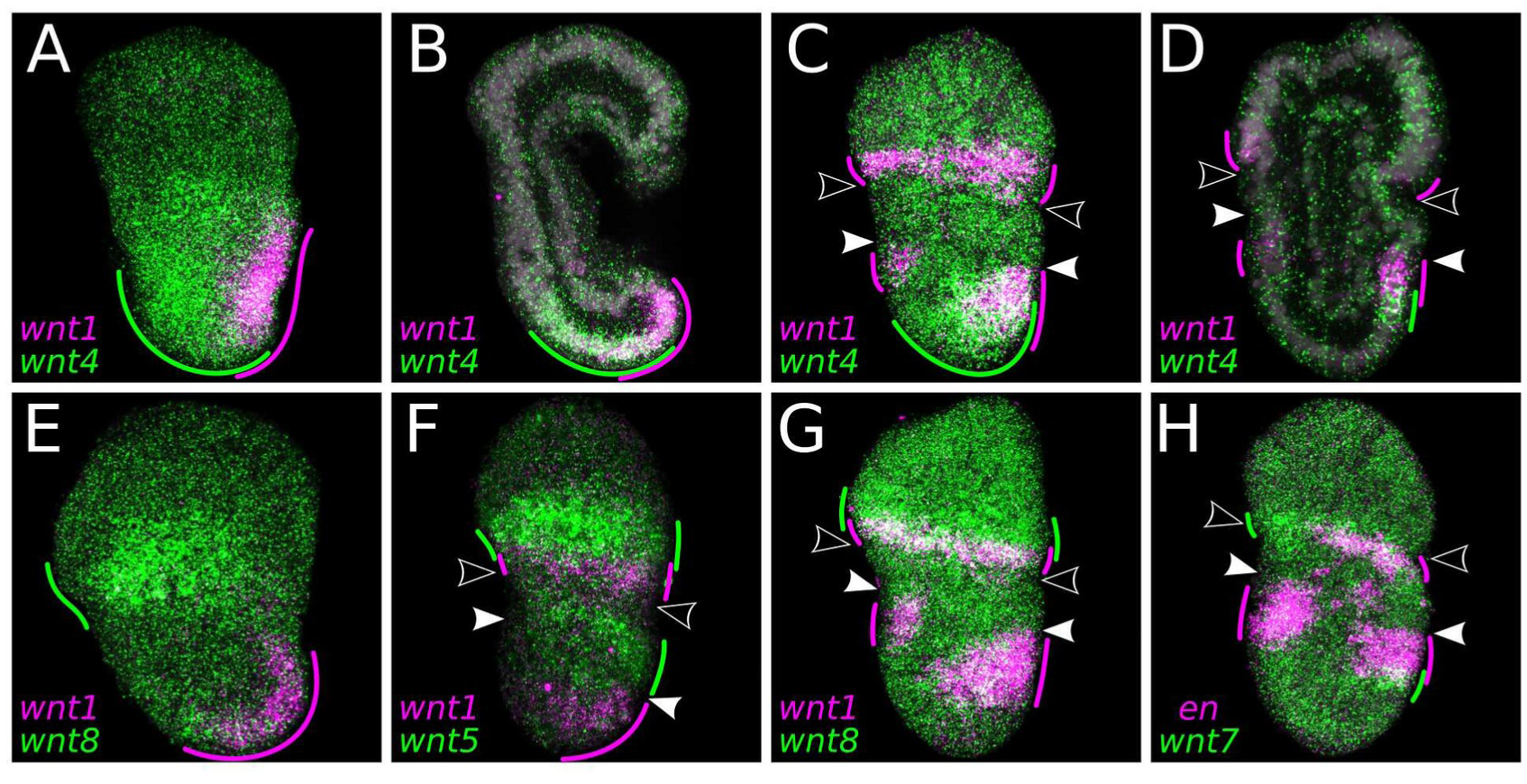
[PNG] Whole-mount double-fluorescent *in situ* hybridization of *Terebratalia transversa wnt* genes. (A–D) Expression of *wnt1* (magenta) and *wnt4* (green) in the mid gastrula (A,B) and late gastrula (C,D). (E,G) Expression of *wnt1* (magenta) and *wnt8* (green) in the mid gastrula (E) and early larva (G). (F) Expression of *wnt1* (magenta) and *wnt5* (green) in the late gastrula. (H) Expression of *engrailed* (magenta) and *wnt7* (green) in the early larva. Green and magenta lines highlight the extension and overlap between domains. Areas in the tissue where the expression overlaps appear in white. Samples oriented with anterior end to the top and ventral to the right (lateral views). Black arrowheads indicate the apical–mantle boundary. White arrowheads demarcate the mantle–pedicle boundary. The panels show representative expression patterns for each sample.

**Additional file 6: Fig. S6:**
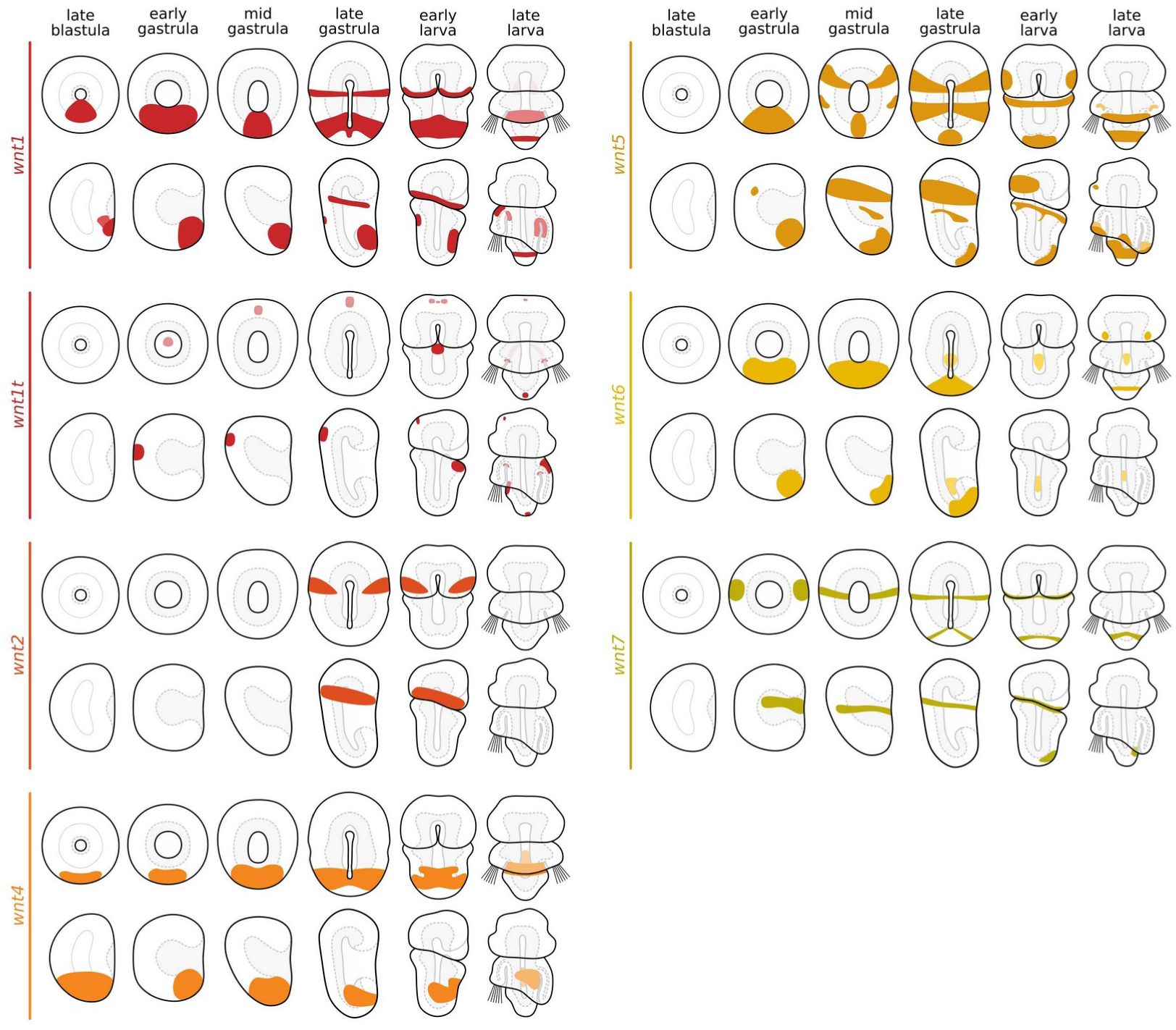
[PDF] Schematic drawings summarizing the expression of *wnt1*, *wnt1t*, *wnt2*, *wnt4*, *wnt5*, *wnt6*, and *wnt7* in *Terebratalia transversa*. For each developmental stage of each gene, a blastoporal/ventral view (top) and a lateral view (bottom) are shown. Faded colors represent expression domains in the mesoderm or endoderm, or in the ectoderm when it is beneath the mantle lobe.

**Additional file 7: Fig. S7:**
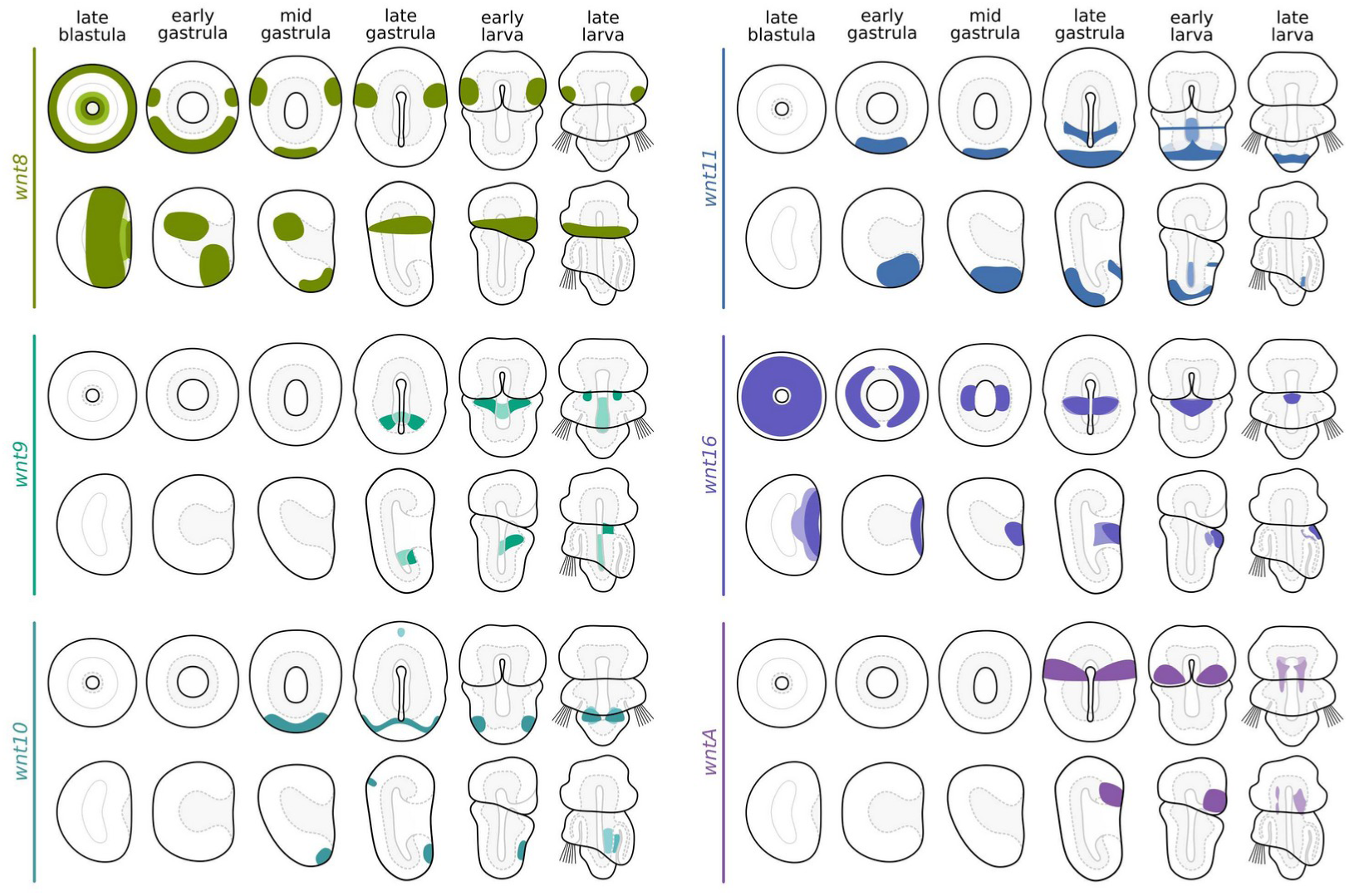
[PDF] Schematic drawings summarizing the expression of *wnt8*, *wnt9*, *wnt10*, *wnt11*, *wnt16*, and *wntA* in *Terebratalia transversa*. For each developmental stage of each gene, a blastoporal/ventral view (top) and a lateral view (bottom) are shown. Faded colors represent expression domains in the mesoderm or endoderm, or in the ectoderm when it is beneath the mantle lobe.

**Additional file 8: Fig. S8:**
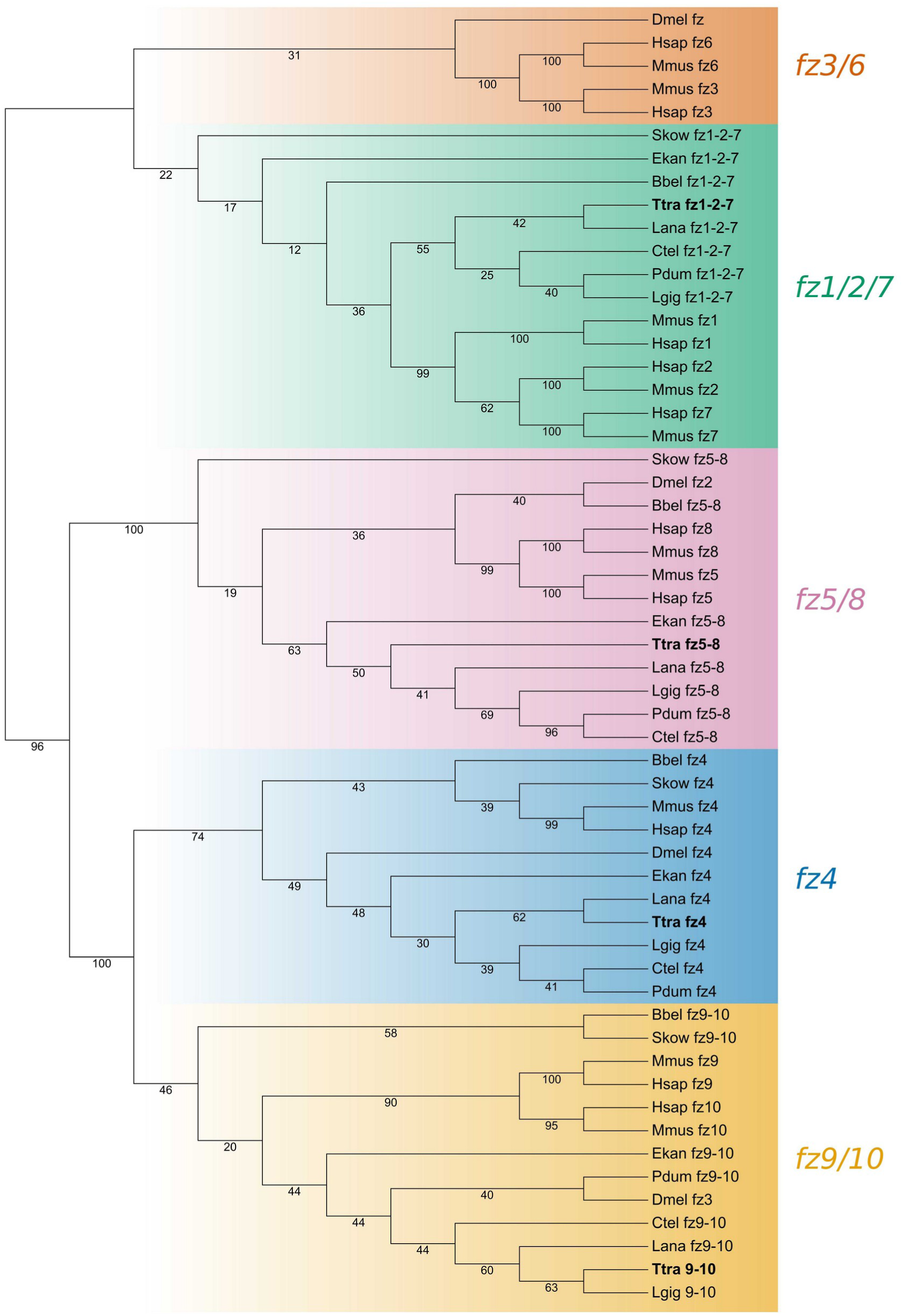
[PDF] Orthology assignment of *Terebratalia transversa* Frizzled proteins. Best-scoring tree of a maximum likelihood phylogenetic analysis using the amino acid sequences of well-annotated Frizzled proteins. The color-coding represents different Frizzled subfamilies and the numbers show the support values of individual branches. *Terebratalia transversa* (Ttra) orthologs are highlighted in bold. The other species are *Branchiostoma belcheri* (Bbel), *Capitella teleta* (Ctel), *Drosophila melanogaster* (Dmel), *Euperipatoides kanangrensis* (Ekan), *Homo sapiens* (Hsap), *Lingula anatina* (Lana), *Lottia gigantea* (Lgig), *Mus musculus* (Mmus), *Platynereis dumerilii* (Pdum), and *Saccoglossus kowalevskii* (Skow).

**Additional file 9: Fig. S9:**
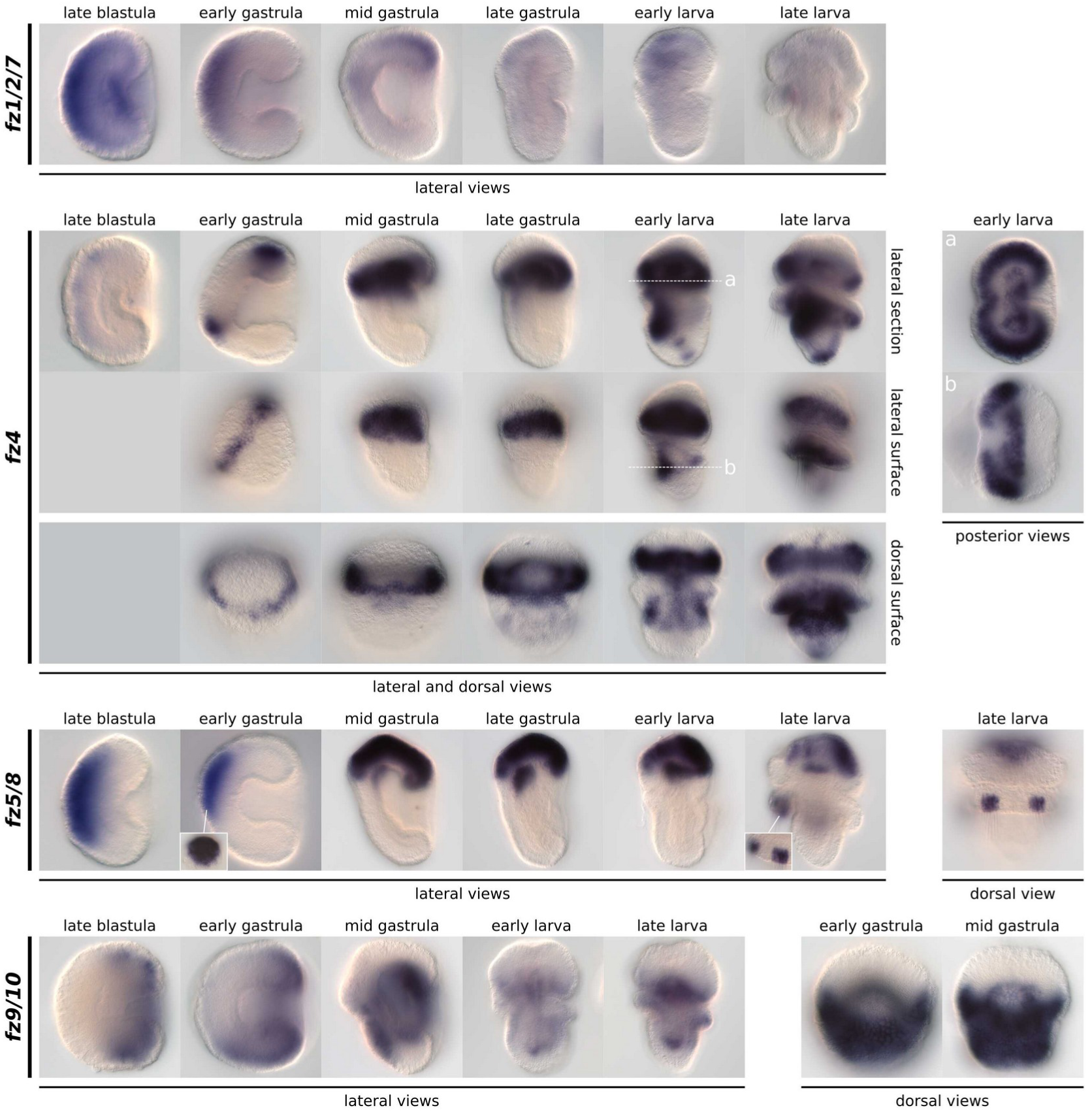
[PNG] Whole-mount colorimetric *in situ* hybridization of *Terebratalia transversa* Frizzled genes. Additional views of *fz1/2/7*, *fz4*, *fz5/8*, and *fz9/10* expression between late blastula and late larva. The panels show representative expression patterns for each sample. The stainings for *fz1/2/7* in the samples from early gastrula to late larva are underdeveloped. Dashed lines indicate the position of the optical section shown in adjacent panels.

**Additional file 10: Fig. S10:**
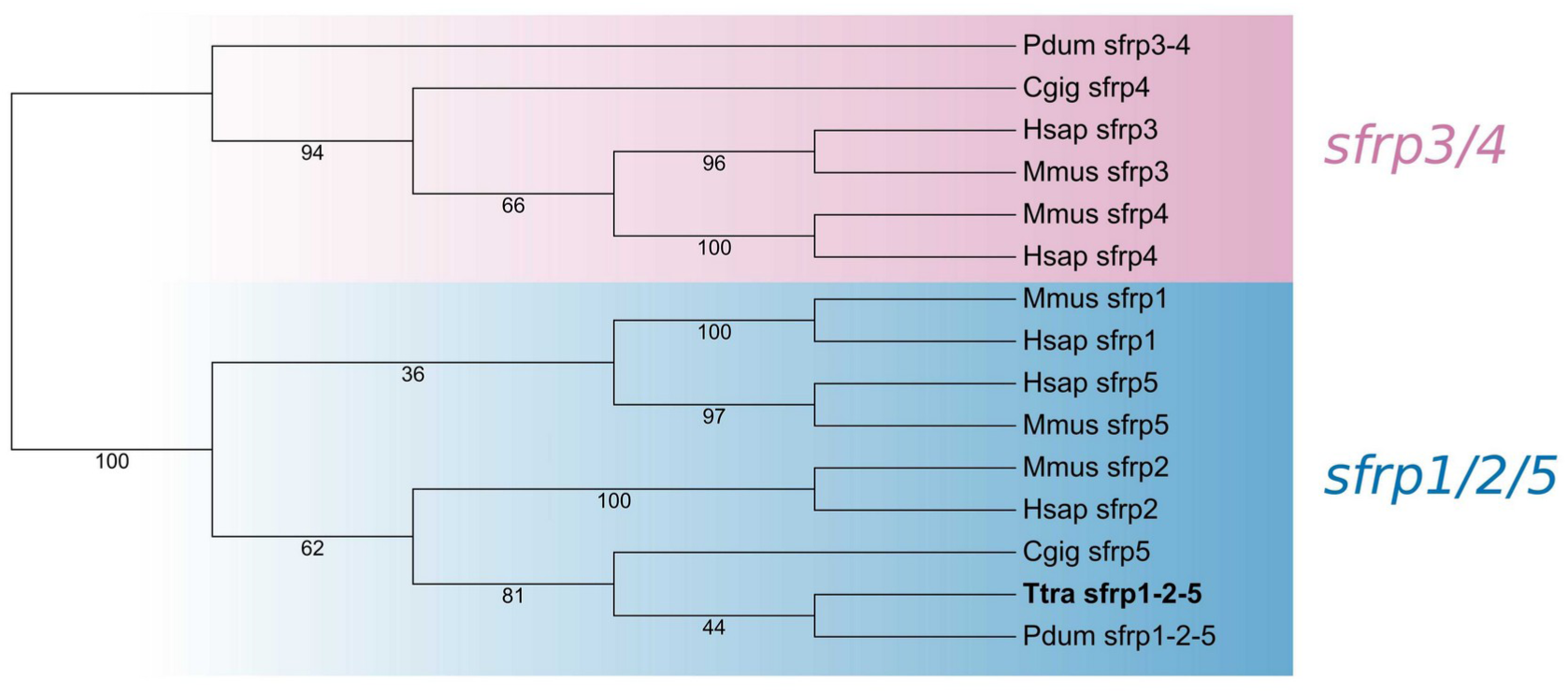
[PDF] Orthology assignment of *Terebratalia transversa* sFRP proteins. Best-scoring tree of a maximum likelihood phylogenetic analysis using the amino acid sequences of sFRP genes. The color-coding represents different sFRP subfamilies and the numbers show the support values of individual branches. *Terebratalia transversa* (Ttra) ortholog is highlighted in bold. The other species are *Homo sapiens* (Hsap), *Crassostrea gigantea* (Cgig), *Mus musculus* (Mmus), and *Platynereis dumerilii* (Pdum).

**Additional file 11: Fig. S11:**
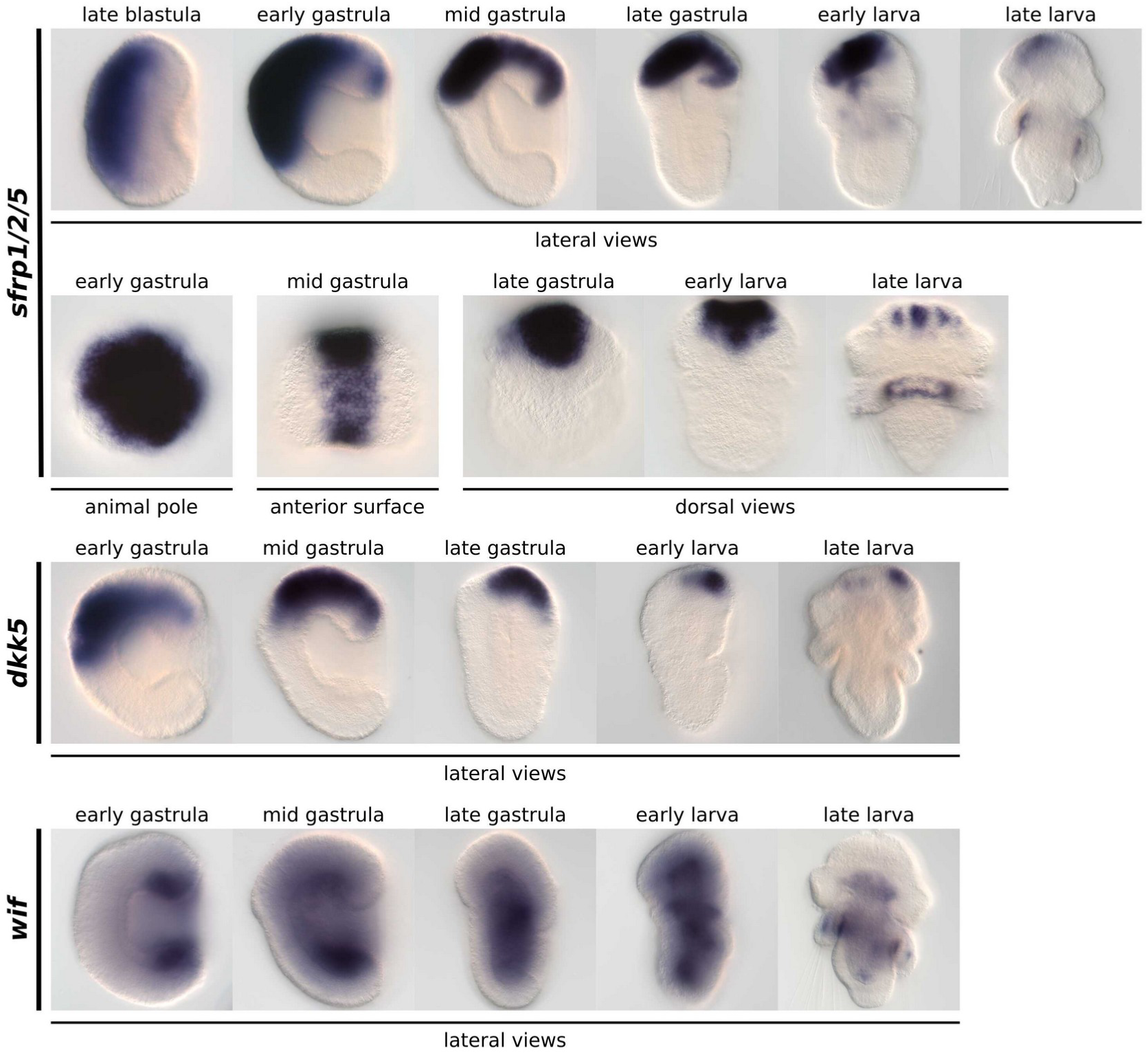
[PNG] Whole-mount colorimetric *in situ* hybridization of *Terebratalia transversa* Wnt antagonists. Additional views of Wnt antagonists expression between late blastula and late larva. The panels show representative expression patterns for each sample.

**Additional file 12: Fig. S12:**
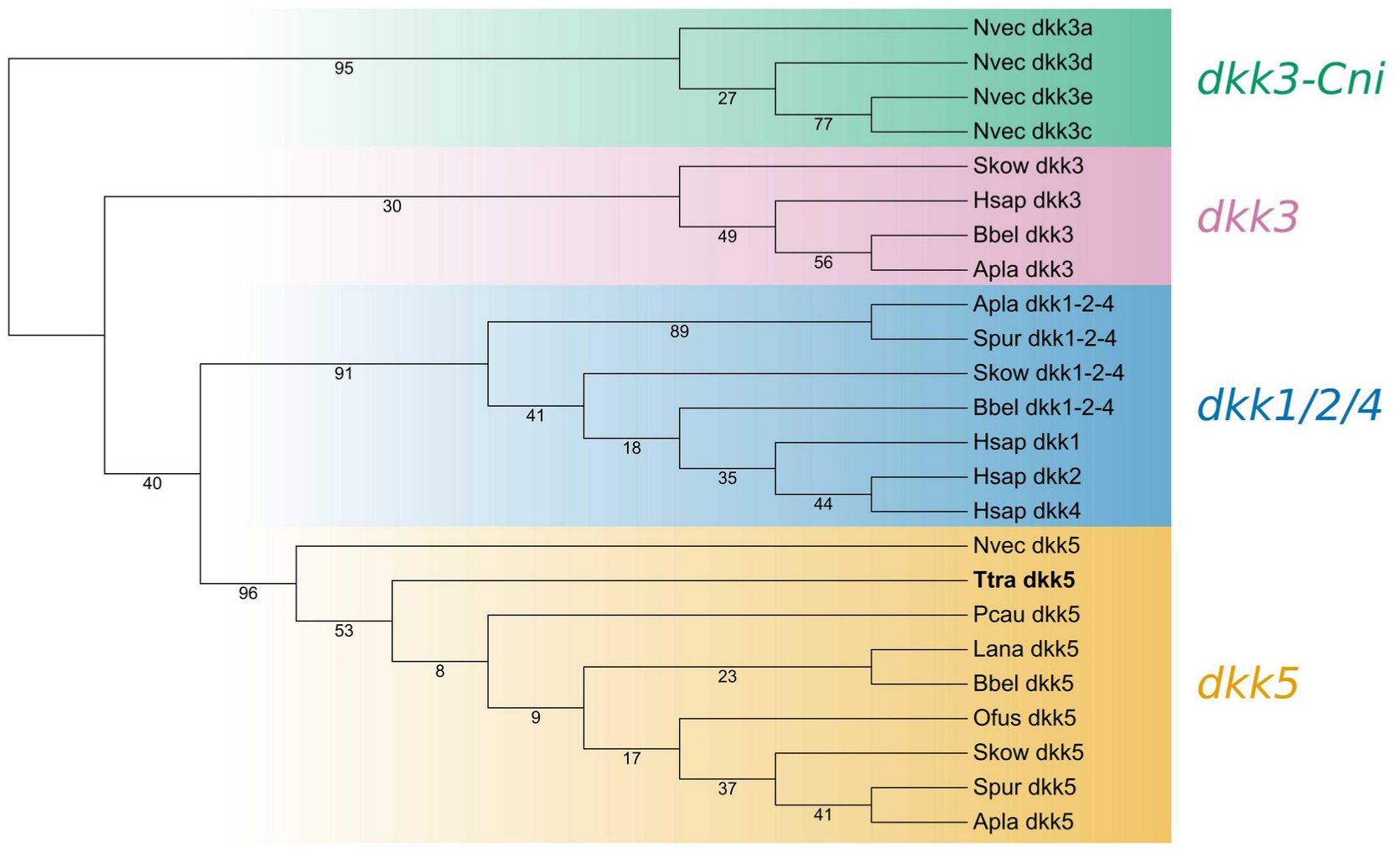
[PDF] Orthology assignment of *Terebratalia transversa* Dkk proteins. Best-scoring tree of a maximum likelihood phylogenetic analysis using the amino acid sequences of Dkk from diverse metazoans. Color-coding represents different Dkk subfamilies. Numbers show support values of individual branches. *Terebratalia transversa* (Ttra) ortholog, highlighted in bold, groups with a previously unidentified Dkk subfamily, in addition to *dkk3* and *dkk1/2/4*, which we named *dkk5*. Non-vertebrate deuterostomes such as the hemichordate *Saccoglossus kowalevskii* (Skow), the echinoderm *Acanthaster planci* (Apla), and the cephalochordate *Branchiostoma belcheri* (Bbel), have an ortholog of each Dkk family. Vertebrates lost *dkk5*. Protostomes lost *dkk1/2/4* and *dkk3* early on, but retained *dkk5* in some lineages such as *T. transversa*, *Priapulus caudatus* (Pcau), and *Owenia fusiformis* (Ofus). Cnidarians expanded *dkk3* but lost *dkk1/2/4*. Overall, this suggests *dkk1/2/4*, *dkk3*, and *dkk5* were the ancestral subfamilies in the Cnidaria–bilaterian branch. The other species are *Homo sapiens* (Hsap), *Lingula anatina* (Lana), *Nematostella vectensis* (Nvec), and *Strongylocentrotus purpuratus* (Spur).

**Additional file 13: Fig. S13:**
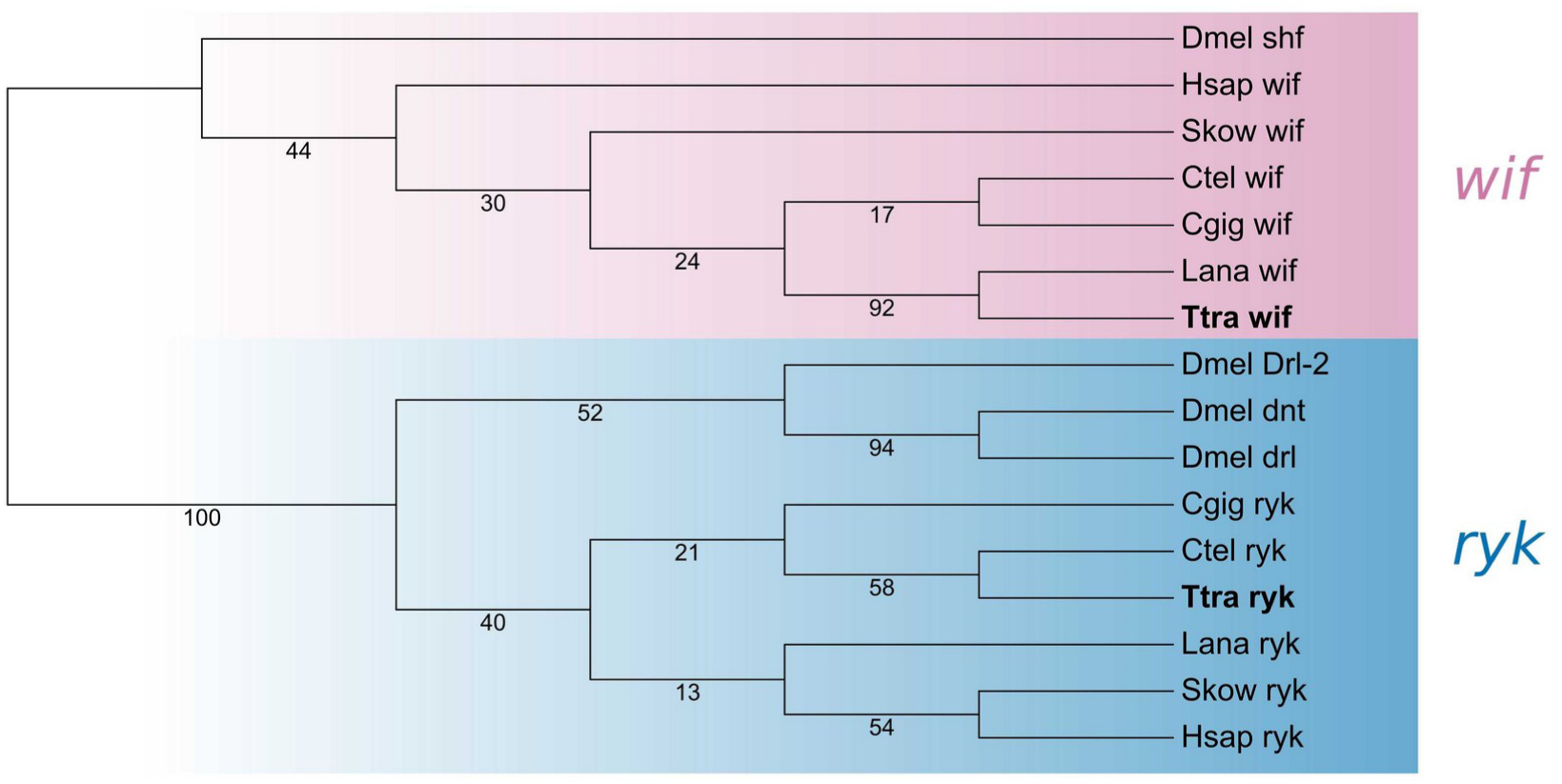
[PDF] Orthology assignment of *Terebratalia transversa* Wif proteins. Best-scoring tree of a maximum likelihood phylogenetic analysis using the amino acid sequences of known Wif proteins (Wnt inhibitory factor). As an outgroup, we used the tyrosine-protein kinase Ryk which also has a WIF domain. The color-coding represents Wif and Ryk families. Numbers show the support values of individual branches. *Terebratalia transversa* (Ttra) orthologs are highlighted in bold. The other species are *Capitella teleta* (Ctel), *Crassostrea gigantea* (Cgig), *Drosophila melanogaster* (Dmel), *Homo sapiens* (Hsap), *Lingula anatina* (Lana), and *Saccoglossus kowalevskii* (Skow).

**Additional file 14: Fig. S14:**
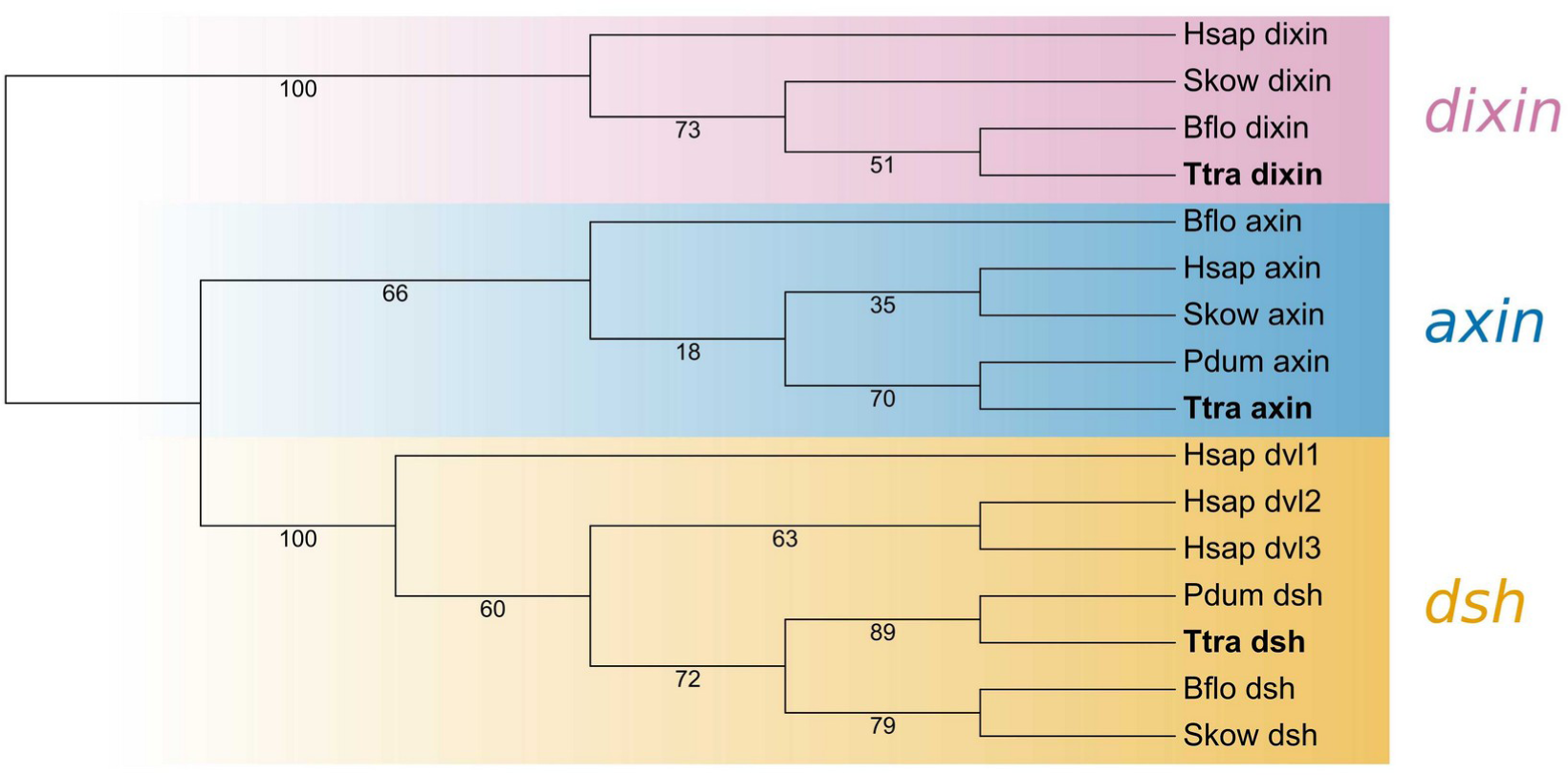
[PDF] Orthology assignment of *Terebratalia transversa* Dsh proteins. Best-scoring tree of a maximum likelihood phylogenetic analysis using the amino acid sequences of known Dsh, Axin, and Dixin proteins. The three belong to the DIX domain superfamily. Each family is color-coded, and the numbers show support values of individual branches. *Terebratalia transversa* (Ttra) orthologs are highlighted in bold. The other species are *Branchiostoma floridae* (Bflo), *Homo sapiens* (Hsap), *Platynereis dumerilii* (Pdum), and *Saccoglossus kowalevskii* (Skow).

**Additional file 15: Fig. S15:**
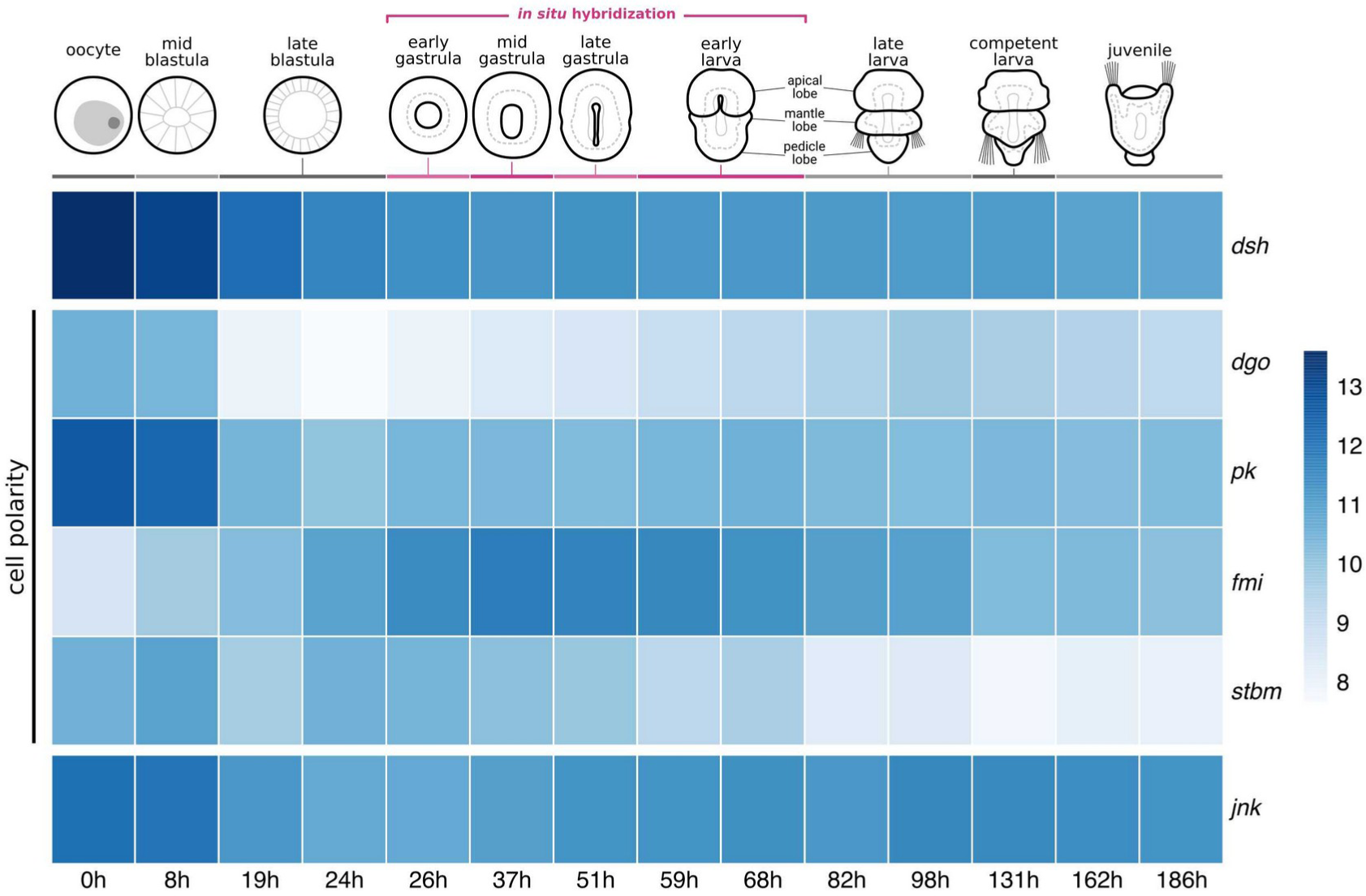
[PDF] Expression of Wnt/PCP pathway during *Terebratalia transversa* development. The heatmap represents the log-normalized transcript counts for *dsh*, *dgo*, *pk*, *fmi*, *stbm*, and *jnk* calculated from stage-specific RNASeq data. Each cell shows the average value between two replicates. The illustrations depict *T. transversa* developmental stages from the oocyte until the post-metamorphic juvenile. The stages we analyzed using *in situ* hybridization (early gastrula to late larva) are highlighted in magenta.

**Additional file 16: Fig. S16:**
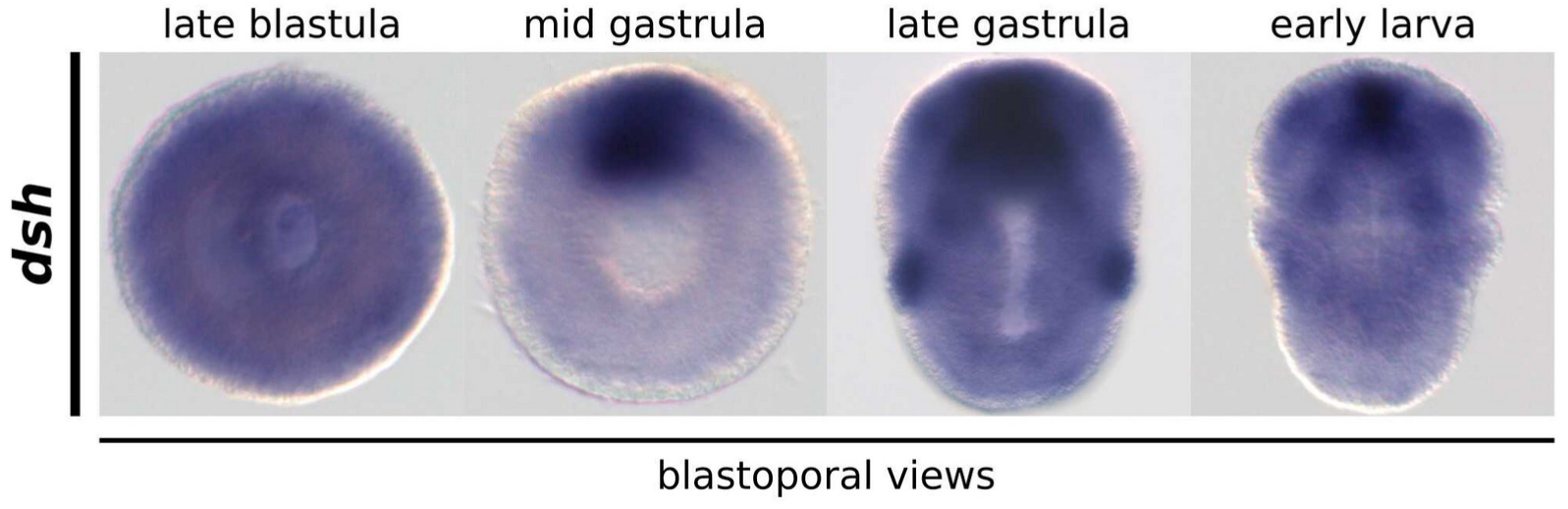
[PNG] Over-developed whole-mount colorimetric *in situ* hybridization of *Terebratalia transversa dsh* gene. The longer reaction time reveals that *dsh* transcripts are ubiquitously expressed in most embryonic tissues.

**Additional file 17: Fig. S17:**
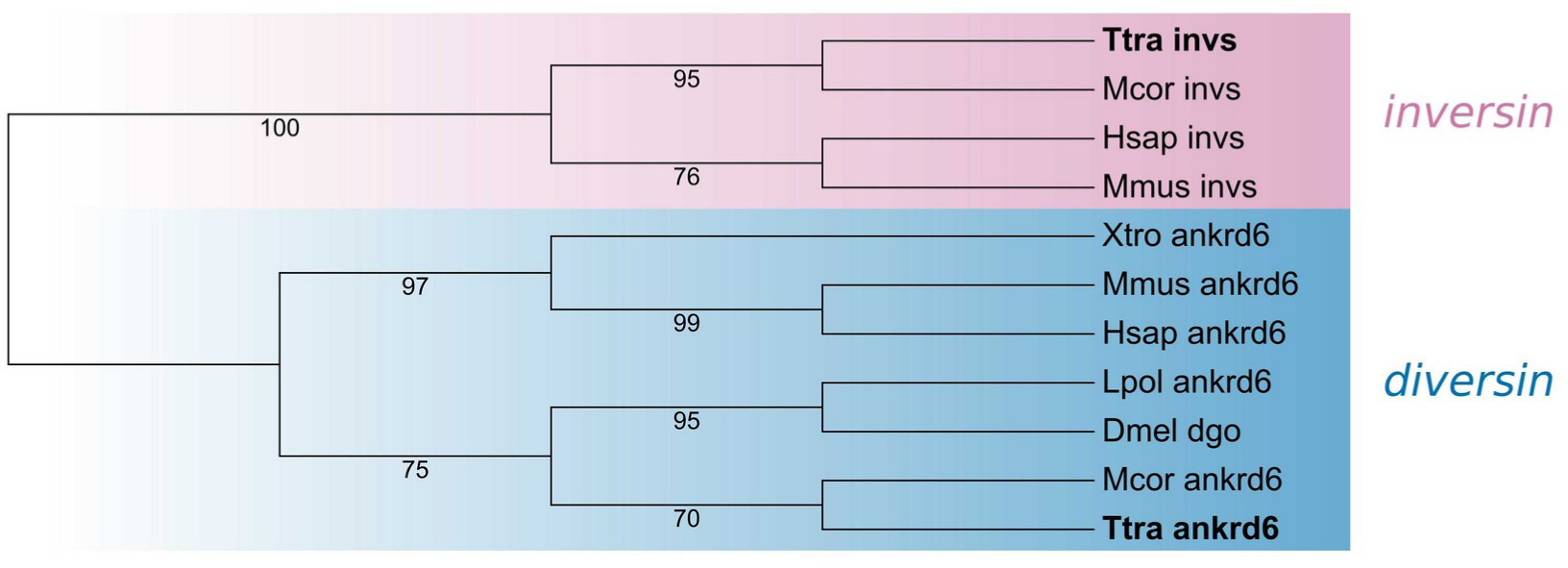
[PDF] Orthology assignment of *Terebratalia transversa* Dgo proteins. Best-scoring tree of a maximum likelihood phylogenetic analysis using the amino acid sequences of Dgo (ANKRD6 or Diversin). We used Inversin proteins as an outgroup since they also have ankyrin repeats. *Terebratalia transversa* (Ttra) orthologs are highlighted in bold. The other species are *Drosophila melanogaster* (Dmel), *Homo sapiens* (Hsap), *Limulus polyphemus* (Lpol), *Mus musculus* (Mmus), *Mytilus coruscus* (Mcor), and *Xenopus tropicalis* (Xtro).

**Additional file 18: Fig. S18:**
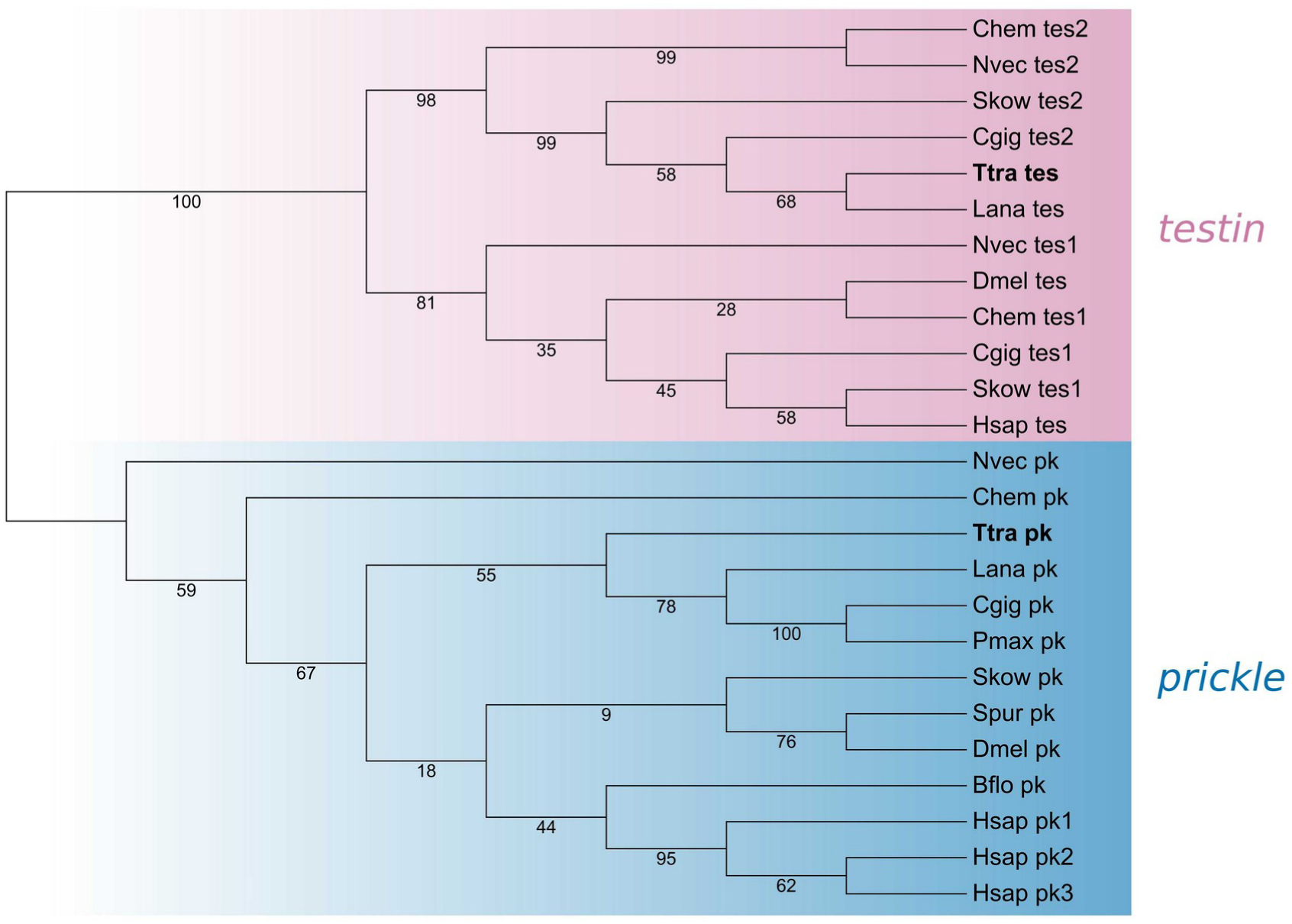
[PDF] Orthology assignment of *Terebratalia transversa* Pk proteins. Best-scoring tree of a maximum likelihood phylogenetic analysis using the amino acid sequences of Pk from diverse metazoans. As an outgroup, we used Testin, a related protein which also contains a LIM and a PET domain. *Terebratalia transversa* (Ttra) orthologs are highlighted in bold. The other species are *Clytia hemisphaerica* (Chem), *Crassostrea gigantea* (Cgig), *Drosophila melanogaster* (Dmel), *Homo sapiens* (Hsap), *Lingula anatina* (Lana), *Nematostella vectensis* (Nvec), *Pecten maximus* (Pmax), and *Saccoglossus kowalevskii* (Skow).

**Additional file 19: Fig. S19:**
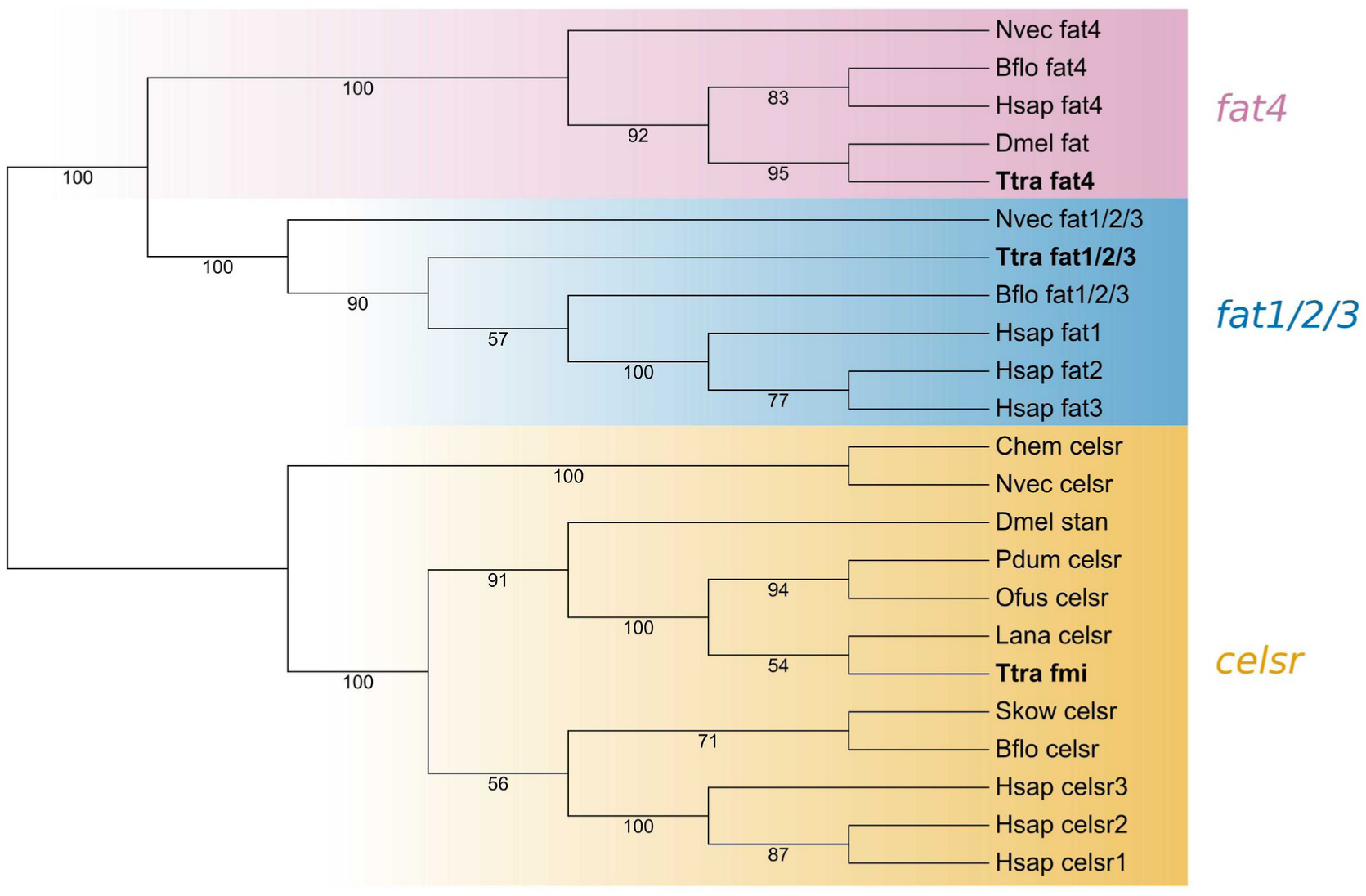
[PDF] Orthology assignment of *Terebratalia transversa* Fmi proteins. Best-scoring tree of a maximum likelihood phylogenetic analysis using the amino acid sequences of Fmi. As outgroups, we used the related Fat family protocadherins which also contain cadherin and laminin domains. *Terebratalia transversa* (Ttra) orthologs are highlighted in bold. The other species are *Branchiostoma floridae* (Bflo), *Clytia hemisphaerica* (Chem), *Drosophila melanogaster* (Dmel), *Homo sapiens* (Hsap), *Lingula anatina* (Lana), *Nematostella vectensis* (Nvec), *Owenia fusiformis* (Of us), *Platynereis dumerilii* (Pdum), and *Saccoglossus kowalevskii* (Skow).

**Additional file 20: Fig. S20:**
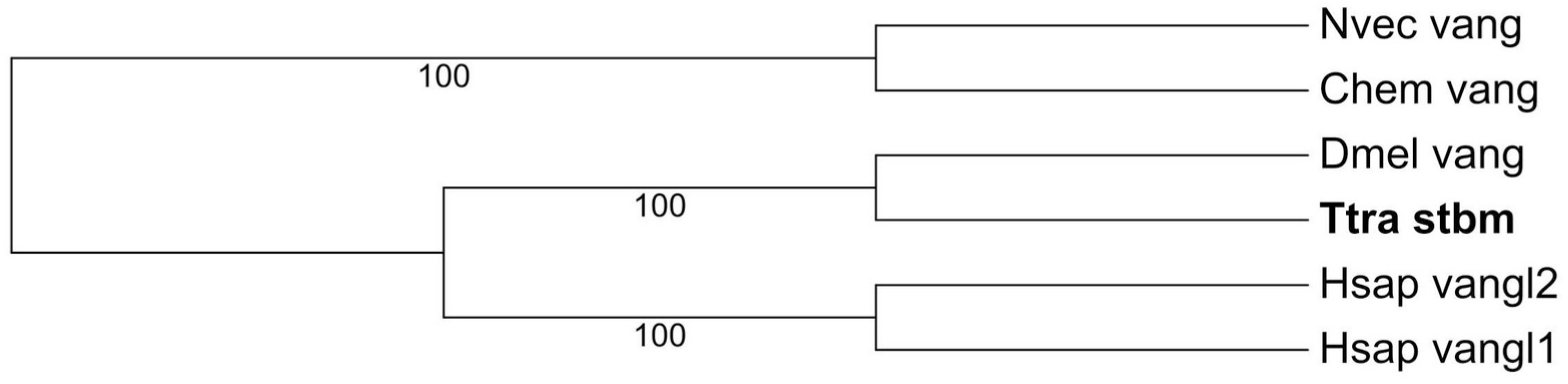
[PDF] Orthology assignment of *Terebratalia transversa* Stbm proteins. Best-scoring tree of a maximum likelihood phylogenetic analysis using the amino acid sequences of Stbm from selected metazoans. *Terebratalia transversa* (Ttra) ortholog is highlighted in bold. The other species are *Clytia hemisphaerica* (Chem), *Drosophila melanogaster* (Dmel), *Homo sapiens* (Hsap), and *Nematostella vectensis* (Nvec).

**Additional file 21: Fig. S21:**
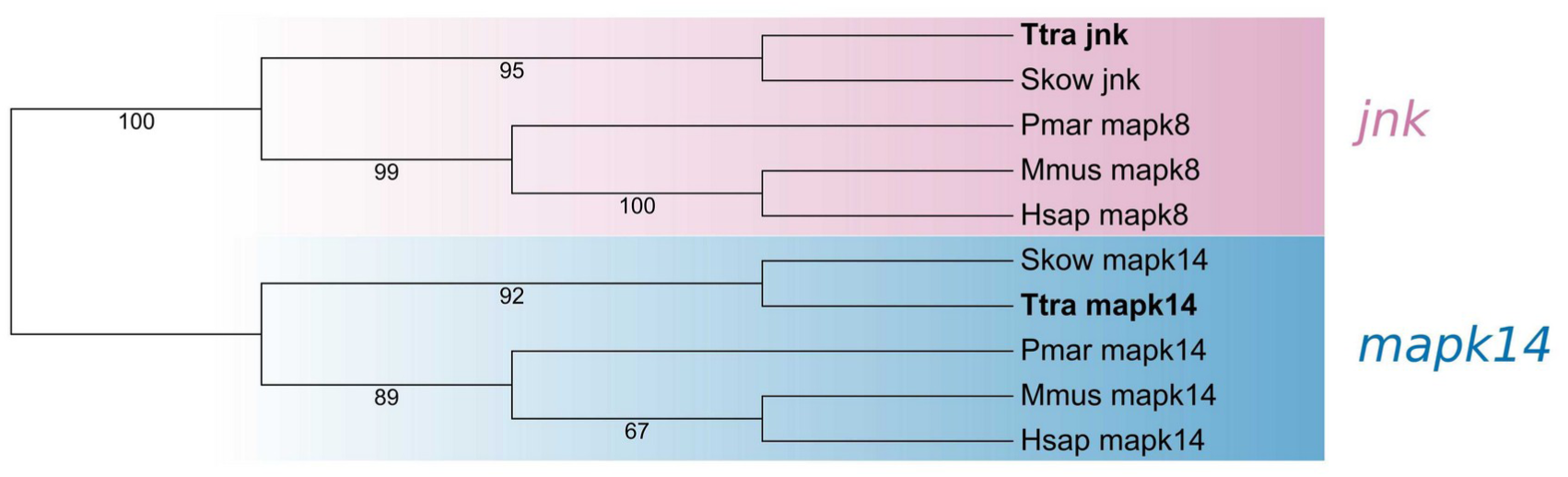
[PDF] Orthology assignment of *Terebratalia transversa* Jnk proteins. Best-scoring tree of a maximum likelihood phylogenetic analysis using the amino acid sequences of Jnk. As outgroup, we used the related protein Mapk14. *Terebratalia transversa* (Ttra) orthologs are highlighted in bold. The other species are *Homo sapiens* (Hsap), *Mus musculus* (Mmus), *Petromyzon marinus* (Pmar), and *Saccoglossus kowalevskii* (Skow).

**Additional file 22: Fig. S22:**
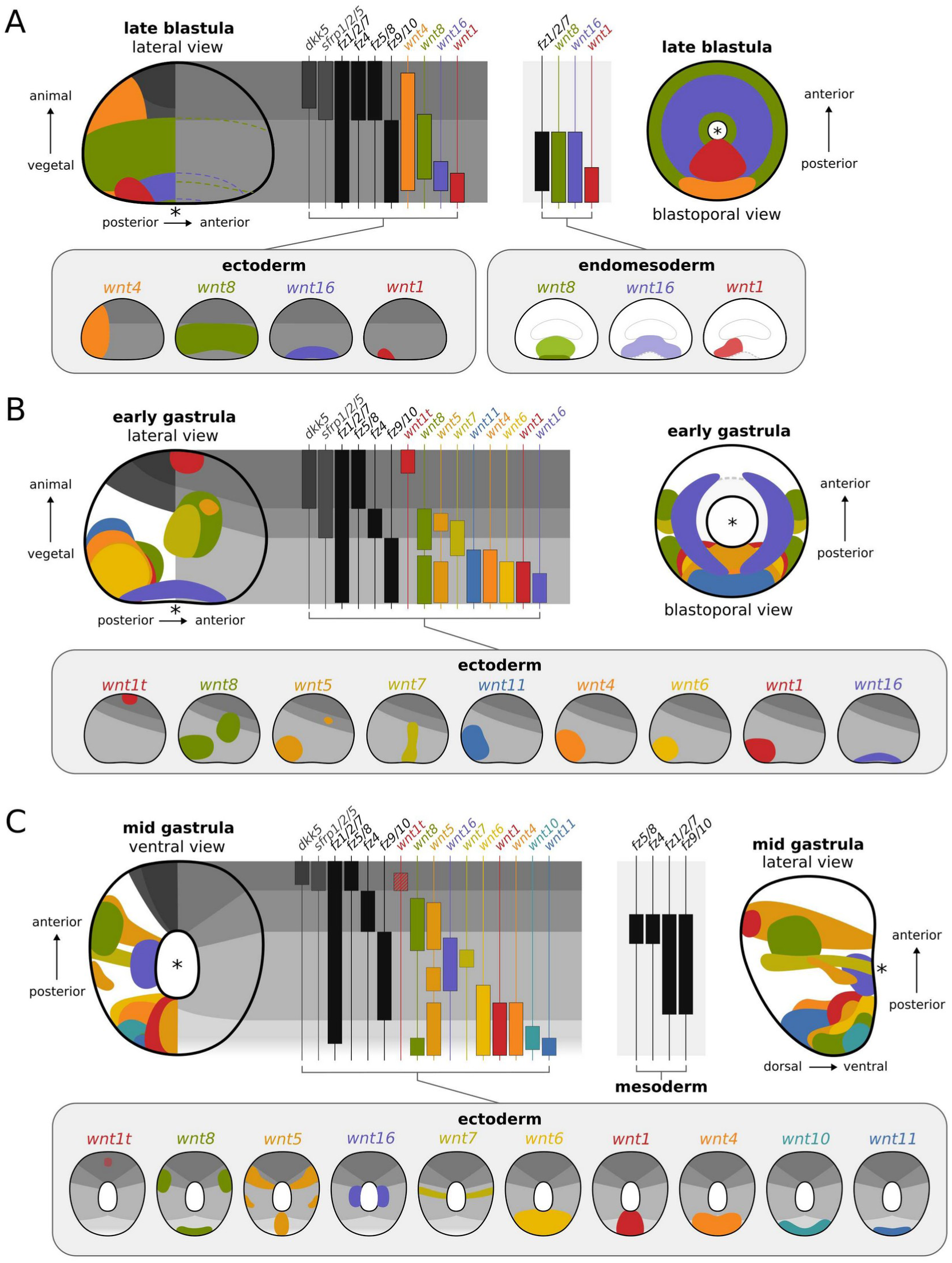
[PDF] Summary of *Terebratalia transversa* Wnt signaling landscape during early gastrulation. Schematic drawings of Wnt genes colored by subfamilies, Frizzled genes by lighter shades of gray, and antagonists by darker shades of gray. The spatial localization of expression domains is superimposed on the embryo (left) and projected to highlight the individualized Wnt genes within the different transcriptional subregions grouped by germ layer (right). The gray boxes show the pattern of individual genes mapped to the embryo for clearer visualization of overlapping domains. (A) Late blastula in lateral and blastoporal views. (B) Early gastrula in lateral and blastoporal views. (C) Mid gastrula in blastoporal/ventral and lateral views.

**Additional file 23: Fig. S23:**
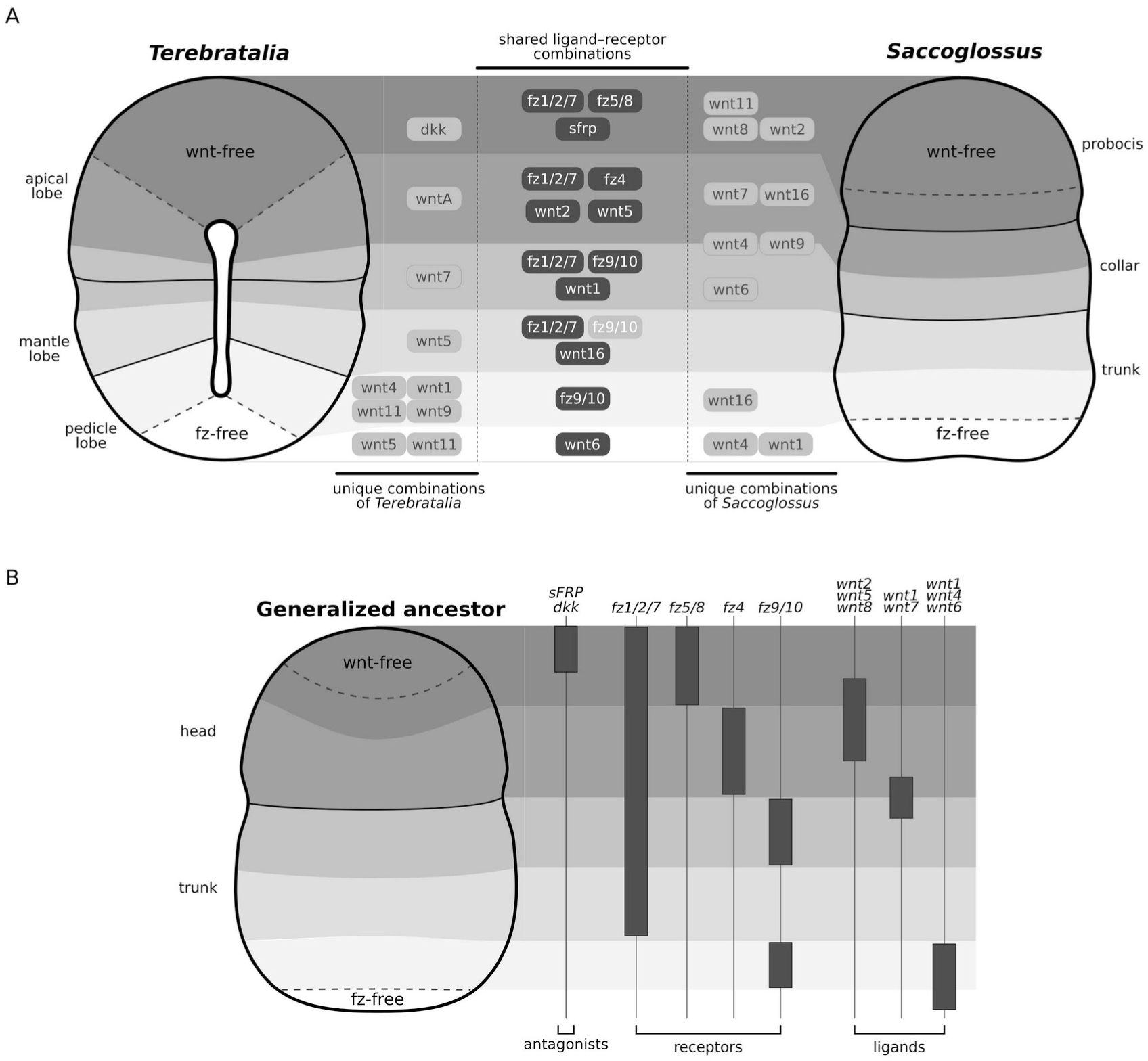
[PDF] Wnt signaling ligand–receptor contexts compared between *Terebratalia transversa* and *Saccoglossus kowalevskii*. (A) Detailed comparison of shared and unique combinations of Wnt signaling components in brachiopod and hemichordate embryos. Solid lines represent morphological boundaries for the apical, mantle, and pedicle lobes, and dashed lines represent boundaries between transcriptional subregions. (B) Generalized ancestor showing the conserved Wnt subregions along the anteroposterior axis of *T. transversa* and *S. kowalevskii*.

**Additional file 24: Fig. S24:**
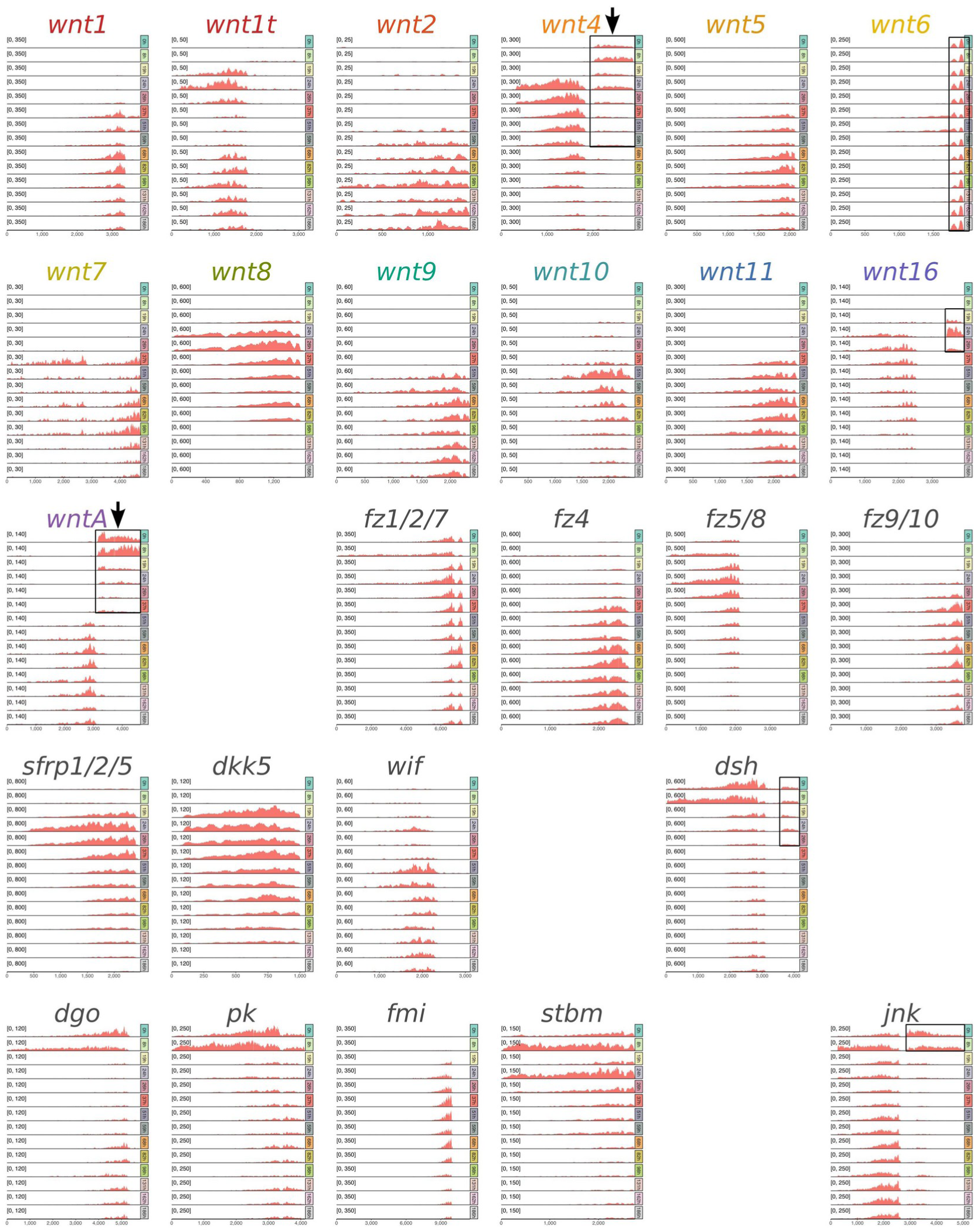
[PNG] Read coverage of the stage-specific transcriptome mapped to the transcripts of *Terebratalia transversa* Wnt signaling components. Each gene shows the read coverage of one replicate along the transcript length for the 14 developmental stages sampled in this study (0–186h). The Y axes are fixed to the maximum observed coverage of a gene (which is different for each gene). The black boxes highlight regions of uneven coverage. Arrows indicate the two cases, *wnt4* and *wntA*, where the uneven coverage caused a bias in the quantification of expression levels. Although *wnt6*, *wnt16*, *dsh*, and *jnk* also show regions of uneven coverage, these reads did not alter the main expression profile of the gene. See the Methods section for more details.

**Additional file 25: Table S1:**
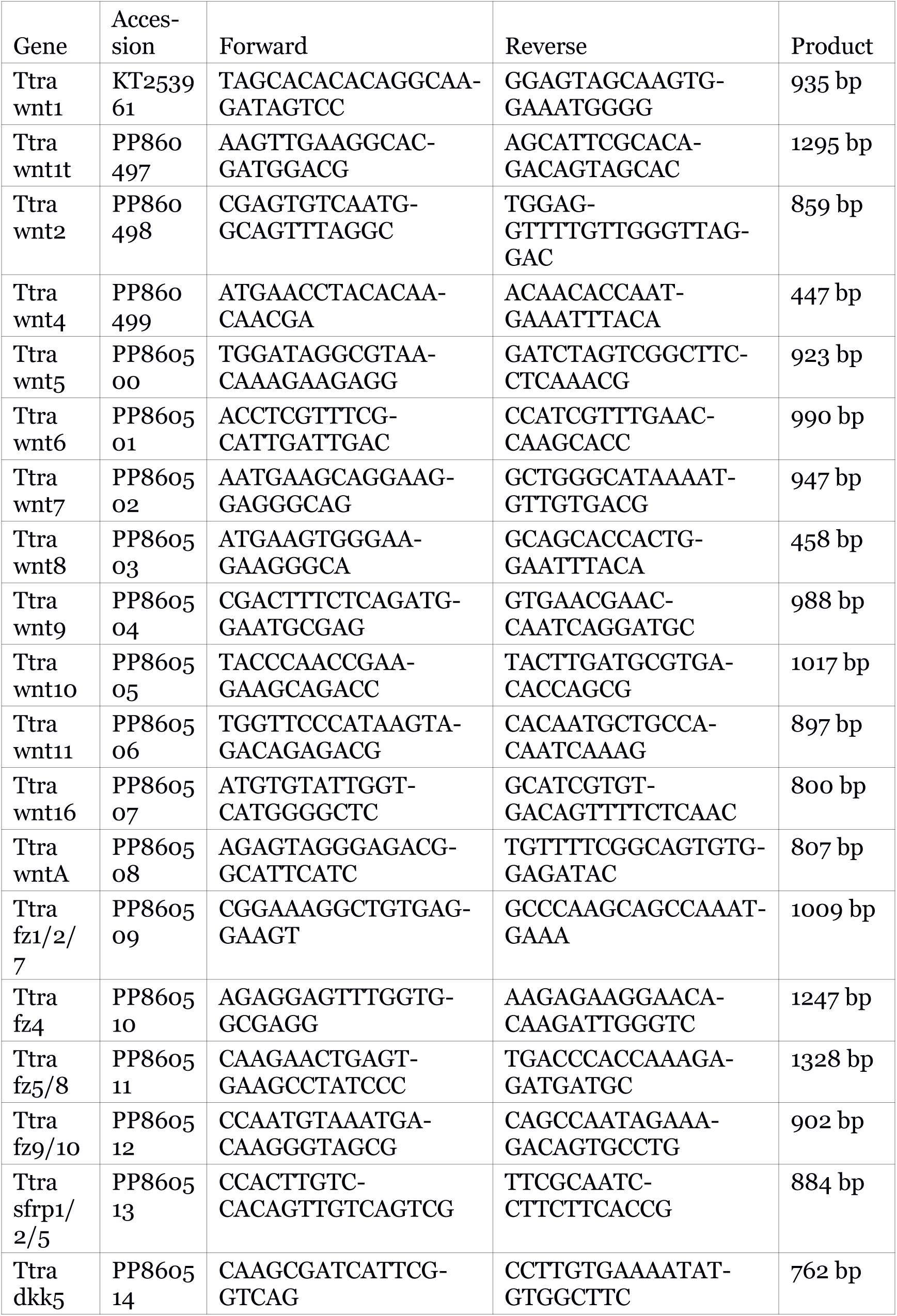
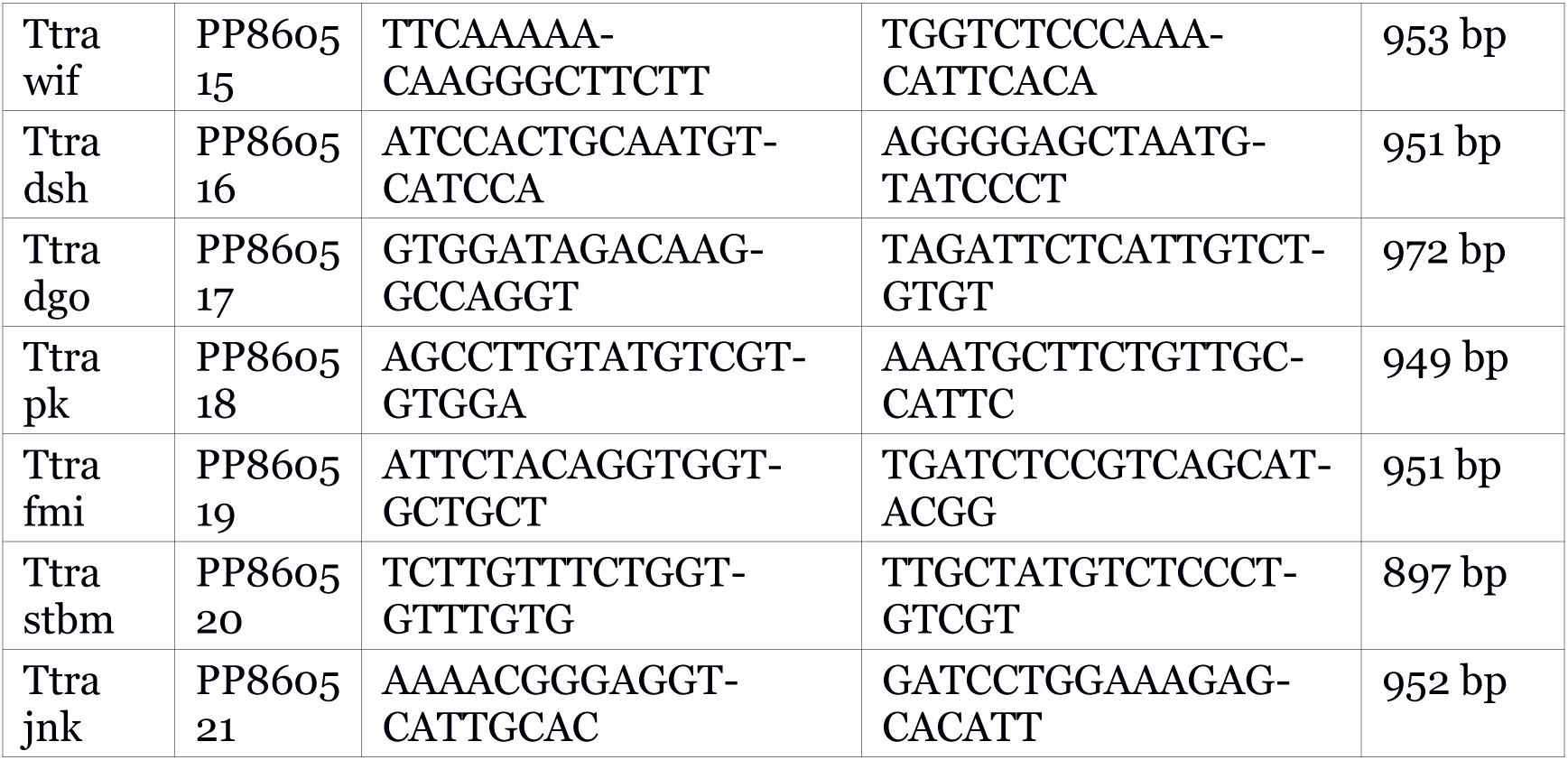
[PDF] Gene accession numbers and primer pairs used for cloning the Wnt signaling components of the brachiopod Terebratalia transversa.

## Notes

https://zenodo.org/doi/10.5281/zenodo.8312022

### Competing Interest Statement

The authors have declared no competing interest.

### Summary of Updates

We improved the readability of the coverage analysis figure, added citations for the tools used in the coverage analysis, and adjusted the figure referencing and manuscript sections.

